# Mutation signature filtering enables high-fidelity RNA structure probing at all four nucleobases with DMS

**DOI:** 10.1101/2023.04.10.536308

**Authors:** David Mitchell, Jennifer Cotter, Irfana Saleem, Anthony M. Mustoe

## Abstract

Chemical probing experiments have transformed RNA structure analysis, enabling high-throughput measurement of base-pairing in living cells. Dimethyl sulfate (DMS) is one of the most widely used structure probing reagents and has played a prominent role in enabling next-generation single-molecule probing analyses. However, DMS has traditionally only been able to probe adenine and cytosine nucleobases. We previously showed that, using appropriate conditions, DMS can also be used to interrogate base-pairing of uracil and guanines *in vitro* at reduced accuracy. However, DMS remained unable to informatively probe guanines in cells. Here, we develop an improved DMS mutational profiling (MaP) strategy that leverages the unique mutational signature of N^1^-methylguanine DMS modifications to enable robust, high-fidelity structure probing at all four nucleotides, including in cells. Using information theory, we show that four-base DMS reactivities convey greater structural information than comparable two-base DMS and SHAPE probing strategies. Four-base DMS experiments further enable improved direct base-pair detection by single-molecule PAIR analysis, and ultimately support RNA structure modeling at superior accuracy. Four-base DMS probing experiments are easily performed and will broadly facilitate improved RNA structural analysis in living cells.

## INTRODUCTION

RNA molecules fold into complex base-paired secondary and tertiary structures that are critically linked to RNA function (1). One of the oldest and most scalable methods for interrogating RNA structure are chemical probing experiments (2-4). These experiments, which can be performed both *in vitro* and in cells, use small molecule chemical reagents to selectively modify flexible nucleotides, yielding per-nucleotide reactivity measurements that report on local RNA structure. Probing data are useful as stand-alone measurements, and can also be used to guide structure modeling algorithms to reconstruct global RNA structure (5, 6). Recently, we and others have introduced single-molecule chemical probing experiments as an even more powerful strategy for characterizing complex RNA systems (7). By measuring correlated modification events, single molecule probing experiments enable direct measurement of secondary base-pairing (8, 9), tertiary interactions (10), and ensembles comprising multiple structural states (11-14). Both traditional and single-molecule experiments have provided critical insights into RNA biology (15-22), and increasingly serve as foundational technologies for RNA functional characterization and therapeutic development (19,23-26).

Numerous strategies utilizing diverse chemical reagents have been developed to probe different aspects of RNA structure (4). The oldest and still one of the most used probing reagents is dimethyl sulfate (DMS) (27). DMS methylates the Watson-Crick face of single-stranded adenine (A) and cytosine (C) nucleobases at the N1 and N3 positions, respectively. DMS is readily cell permeable and can be used to achieve high levels of modification, which is essential for single-molecule probing applications.

However, the inability of DMS to probe uracil (U) and guanine (G) bases has represented a key limitation. We recently showed that mildly alkaline conditions enable DMS to effectively probe U structure (8). These conditions also support probing of G structure *in vitro* at lower but still useful accuracy. However, DMS remained unable to probe G structure in cells. Probing G is particularly desirable because of the outsized role G nucleotides play in stabilizing RNA structure. Alternative chemical reagents permit probing of G and U (28-30), and SHAPE (selective 2’-hydroxyl acylation analyzed by primer extension) reagents support probing of all four bases (31-34), but to date these reagents have proven less amenable for single-molecule analyses. Enabling robust DMS probing at all four bases has the potential to broadly improve both per-nucleotide and single-molecule RNA structural analysis.

A seminal advance in chemical probing technology has been the development of mutational profiling (MaP) (10, 35). MaP uses specialized reverse transcription protocols to read through and encode chemical modifications as mutations in cDNA, which can then be measured via high-throughput sequencing. Compared to alternative strategies, MaP permits precise quantitation of even rare modification events. Critically, MaP also permits measurement of multiple modifications per individual molecule, enabling detection of the correlated modification events that underpin single-molecule probing strategies (7). MaP may also enable distinguishment between different chemical modifications that generate distinct mutational signatures, although this application is relatively unexplored.

In recent years, multiple reverse transcriptase enzymes have been adapted for performing DMS-MaP experiments. G- and U-sensitive DMS probing and most single-molecule probing analyses have relied on a relaxed-fidelity Superscript II (SSII) reverse-transcription protocol (8, 10, 11). Newer MaP protocols that use TGIRT-III (TGIRT) and MarathonRT (Marathon) reverse transcriptase have been reported to be more processive and exhibit lower background error rates (36-38). However, these enzymes have not been evaluated for their ability to measure DMS modifications at G and U nucleotides, nor for their ability to support direct base pair detection via single-molecule correlation analysis.

In this work, we sought to evaluate different MaP protocols for their ability to support DMS probing at all four bases. Strikingly, our analyses revealed that reverse transcriptases decode DMS-induced N^1^-methylguanine and N^7^-methylguanine chemical modifications via distinct mutational signatures. Leveraging this discovery, we develop a new strategy that solves prior limitations to enable high-fidelity DMS probing at all four nucleotides in cells. Four-base DMS-MaP conveys more information than other comparable structural probing experiments, facilitates more sensitive single-molecule analysis, and ultimately enables improved RNA structural modeling, representing an all-in-one strategy for complete RNA structural analysis.

## MATERIALS AND METHODS

### Probing of HEK293 cells

*DMS probing:* Human HEK293 cells (ATCC; CRL-1573) were maintained in DMEM supplemented with 10% FBS and 100 U/mL Pen/Strep at 37 °C and 5% CO_2_. ∼1 × 10^6^ cells were seeded in a 6 cm culture dish and grown to ∼75% confluency. Prior to DMS treatment, media was exchanged with 2.8 mL fresh media, followed by addition of 800 µL of 1 M bicine (pH 8.3 at room temperature), 1 M sodium cacodylate (pH 7.2), or nuclease-free water, and equilibrated for 3 min at 22 °C. Cells were modified by adding 400 µL of 1.7 M DMS solution in ethanol (or 100% ethanol for control reactions) and incubating at 37 °C for 6 min. Reactions were quenched by addition of 4 mL ice-cold 20% 2-mercaptoethanol (vol/vol in PBS). Cells were scraped and pelleted by centrifugation at 1000 g for 5 min at 4 °C, followed by RNA extraction with 1 mL TRIzol reagent (ThermoFisher). Genomic DNA was removed by addition of 6 U of TURBO DNase (Ambion) and incubation at 37 °C for 45 min. RNA then was purified (RNA Clean & Concentrator, Zymo), quality assessed by TapeStation analysis (Agilent), and concentration quantified by UV absorbance (NanoDrop, ThermoFisher).

*2A3 probing:* Probing was done as described by Marinus *et al* (34). HEK293 cells were maintained as described above. Prior to 2A3 treatment, media was removed, cells washed with 1x PBS, followed by addition of 1 mL trypsin and 2 min incubation at 37 °C to detach cells from the dish. Trypsin was neutralized by 2 mL of media and cells pelleted by centrifugation at 1000 *g* for 5 min at 22 °C. The cell pellet was resuspended in 90 µL of PBS. 10 µL of 1 M 2A3 (Tocris Bioscience) was added to the cells followed by 15 min incubation at 37 °C with occasional tapping to mix. 2A3 reactions were quenched by addition of 100 mL of ice-cold 20% 2-mercaptoethanol. RNA then was extracted as described above.

### DMS probing of *E. coli* RNA

*Cell-free experiments:* Cell-free DMS probing of *E. coli* K-12 MG1655 total RNA was performed as previously described (8). 148 mL LB was inoculated with 2 mL of an overnight culture and grown at 37 °C until OD_600_ ≈ 0.5. 16.65 mL of 187.5 µg/mL rifampicin was added followed by incubation for 20 min at 37 °C to chase assembly of RNA-protein complexes (39). Cells were pelleted and resuspended in 32 mL of lysis buffer [15 mM Tris HCl (pH 8), 450 mM sucrose, 8 mM EDTA (pH 8)], followed by addition of 1.28 mL of 1 mg/mL lysozyme, and 10 min incubation on ice. Total RNA was extracted by 3x phenol/chloroform/isoamyl alcohol (PCA) extraction, 3x chloroform extraction, and exchange into 1x bicine folding buffer [200 mM bicine (pH 8.3 at room temperature), 200 mM potassium acetate, 5 mM MgCl_2_]. After 10 min equilibration at 37 °C, 1 volume of 1.7 M DMS solution in ethanol (or 1 volume of 100% ethanol for control reactions) was added to 9 volumes of total RNA and reacted for 6 min at 37 °C. Ten volumes of ice-cold 20% 2-mercaptoethanol was added to quench reactions, followed by purification (RNeasy Midi, Qiagen), DNase treatment (TURBO DNase, ThermoFisher), and a final purification (RNeasy Midi, Qiagen). RNA was then quantified for quality (TapeStation, Agilent) and concentration (NanoDrop, ThermoFisher).

*In-cell experiments:* In-cell DMS probing of *E. coli* total RNA was performed as described (8). Cells were grown and treated with rifampicin as described for cell-free experiments, then pelleted by centrifugation at 4000 g for 5 min at 22 °C. Cell pellets were resuspended in 20 mL of 1x bicine folding buffer [200 mM bicine (pH 8.3 at room temperature), 200 mM potassium acetate, 5 mM MgCl_2_]. For experiments using alternative extracellular buffers, pH 8.3 bicine buffer was replaced with 200 mM pH 7.2 sodium cacodylate, 200 mM bicine pH 8, or 200 mM bicine pH 9 (all prepared at room temperature). 4.5 mL of cells was added to 0.5 mL of 7 M DMS in ethanol (or 0.5 mL 100% ethanol for control reactions) and reacted for 6 min at 37 °C. DMS reactions were quenched by addition of 20 mL of ice-cold 20% 2-mercaptoethanol and the cells placed on ice. Cells were pelleted, resuspended in 1 mL of 1mg/mL lysozyme, and incubated on ice for 5 min. Total RNA was extracted using TRIzol reagent, DNase treated (TURBO DNase, Invitrogen), and then purified (RNeasy Midi, Qiagen).

### Reverse Transcription

Four different mutational profiling (MaP) reverse transcription (RT) protocols were evaluated, which we refer to by the RT enzyme used: Superscript II (SSII) (8), MarathonRT (Marathon) (37), TGIRT-III (TGIRT) (36), and evolved HIV RT (40). Gene-specific priming was used for human RNase P and RMRP, and *E. coli* tmRNA, and random priming was used for ribosomal RNAs. RT reactions were purified using magnetic beads (Mag-Bind Total Pure NGS, Omega Bio-Tek). 2A3-probed RMRP and RNase P samples were reverse transcribed using the SSII protocol.

*SSII:* 2 µL of 10 mM dNTPs and 1 µL of either 2 µM specific primer or 200 ng/µL random 9-mer was added to 8 µL of 1-2 µg RNA and incubated at 65 °C for 10 min followed by 4 °C for 2 min. Subsequently, 8 µL of 2.5x SSII MaP buffer [final concentration 50 mM Tris-HCl (pH 8), 75 mM KCl, 6 mM MnCl_2_, 1 M betaine, 10 mM DTT] was added to the solution, followed by 1 µL of SSII (Invitrogen). The reaction was then incubated according to the temperature program: 25 °C for 10 min, 42 °C for 90 min, 10x[50 °C for 2 min, 42 °C for 2 min], 72 °C for 10 min.

*Marathon:* 2 µL of 10 mM dNTPs and 1 µL of either 2 µM specific primer or 200 ng/µL random 9-mer was added to 2.8 µL of 1-2 µg extracted RNA and incubated at 65 °C for 10 min followed by 4 °C for 2 min. Afterward, 12.2 µL of Marathon MaP buffer [final concentration 50 mM Tris-HCl (pH 8.3), 200 mM KCl, 5 mM DTT, 1 mM MnCl_2_, 20% glycerol] and 2 µL of Marathon (Kerafast) were added to the solution. The solution then was incubated at 42 °C for 3 h followed by 95 °C for 1 min.

*TGIRT:* 2 µL of 10 mM dNTPs and 1 µL of either 2 µM specific primer or 200 ng/µL random 9-mer was added to 8 µL of 1-2 µg RNA and incubated at 65 °C for 10 min followed by 4 °C for 2 min. 8 µL of 5X TGIRT-III RT buffer [final concentration 50 mM Tris–HCl (pH 8.3), 75 mM KCl, and 3 mM MgCl_2_] and 1 µL TGIRT-III (InGex) were added to the solution, followed by incubation at 60 °C for 2 h.

*eHIV:* 1 µL of 10 mM dNTPs and 1 µL of 5 µM specific primer was added to 7 µL containing 2 µg of RNA and incubated at 70 °C for 2 min followed by 4 °C for 2 min. 9 µL of 5X eHIV RT buffer [final concentration 200 mM Tris-HCl (pH 8.3), 400 mM KCl, 20 mM MgCl_2_] and 2 µL of eHIV enzyme were added to the solution. The reaction was then incubated at 42 °C for 3 h followed by 95 °C for 1 min. The eHIV enzyme was expressed and purified from pET30-RT1306 (gift from Bryan Dickinson; Addgene plasmid # 131521) following published protocols (40).

### Library Preparation

*Small RNAs:* Libraries were prepared using a two-step PCR strategy (6, 8). 1 µL of cDNA was input into PCR1 using the following temperature cycles: 98 °C for 30 s, 18 cycles of [98 °C for 10 s, 60 °C for 30 s, 72 °C for 20 s], and 72 °C for 2 min. PCR1 products were purified (Mag-Bind Total Pure NGS, Omega Bio-Tek) using a 0.7x bead ratio. 1 ng of PCR1 product was used as input for PCR2 using the following temperature cycles: 98 °C for 30 s, then 12 cycles of [98 °C for 10 s, 66 °C for 30 s, 72 °C for 20 s], and 72 °C for 2 min. PCR2 products were purified using a 0.7x bead ratio. Libraries were sequenced on an Illumina MiSeq instrument using either 2×250 (v2 chemistry) or 2×300 (v3 chemistry) paired-end sequencing.

*rRNAs:* Libraries from randomly primed total RNA were prepared using the xGen NGS RNA (Integrated DNA Technologies) kit for HEK293 and the Nextera XT (Illumina) kit for *E. coli* cells. cDNAs were converted to double stranded DNA (dsDNA) by NEBNext second-strand synthesis module (New England Biolabs) using 2 h incubation at 16 °C. dsDNA was purified and size selected using magnetic beads (Mag-Bind Total Pure NGS, Omega Bio-Tek) using a 0.65x bead ratio. Libraries were then generated following the Nextera XT manufacturer protocol, followed by purification and size-selection by magnetic beads (Mag-Bind Total Pure NGS, Omega Bio-Tek) using a 0.56x bead ratio. Libraries were sequenced on an Illumina MiSeq instrument using 2×300 paired-end sequencing (v3 chemistry).

### Mutation signature analysis

RMRP and RNase P probing data were initially processed using *ShapeMapper* 2.1.5 with the *--output-counted-mutations* flag to tabulate mutation types observed at each sequence position. Area under the receiver operating characteristic curves (AUROC) were calculated with *Scikit-Learn* (0.24.1) in Python using background-subtracted mutation rates with pairing status of each position derived from known reference structures (41-45). To identify the G mutation signature filter, AUROC was calculated for all possible combinations of mutation types, which revealed that including only G-to-C and G-to-T substitutions yielded the highest AUROC for all enzymes.

### ShapeMapper 2.2

Building on our mutation signature analysis, we incorporated several new features into *ShapeMapper* to automate four-base DMS-MaP processing. This new version of *ShapeMapper* (v2.2) is available for download at https://github.com/Weeks-UNC/shapemapper2. DMS-specific processing is invoked using the “--dms” flag.

*Mutation signature filtering:* G-to-A single-nucleotide mismatches, G multi-nucleotide mismatches, and insertions and deletions at all nucleotides are ignored (set internally to “no data”).

*DMS reactivity normalization:* Because four-base DMS modification rates vary significantly based on nucleotide identity, reactivities are normalized on a nucleotide-specific basis. The DMS modification rate is calculated as the difference between the modified and untreated mutation rates, or simply as the modified mutation rate if no untreated sample is provided. Normalization factors for each nucleotide type *n* are computed as

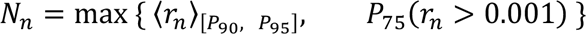

where 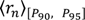 denotes the mean of 90^th^–95^th^ percentile modification rates, *P*_75_(*r_n_* > 0.001) denotes the 75^th^ percentile of modification rates >0.001. This scheme is more robust for RNAs such as the ribosome where most nucleotides are unreactive. Final normalized reactivities are then obtained by dividing the modification rate by the nucleotide-specific normalization factor. The normalized reactivities are output directly as text files with the suffix *.dms*.

### Final data processing

Four-base DMS probing data were processed using *ShapeMapper* 2.2 with the *--dms* and *--output-parsed-mutations* options. Amplicon libraries from small RNAs were processed using the *--amplicon* option, and total RNA (rRNA) libraries were processed using the *--random-primer-len 9* flag. Unfiltered (standard DMS) and 2A3 data were processed using *ShapeMapper* 2.2 without the *--dms* flag.

### Expected Structural Information

Inspired by metrics for quantifying sequence information content (46), we developed the Expected Structural Information (ESI) metric to intuitively quantify the total information provided by a probing experiment. Each nucleotide can adopt two possible structural states: base-paired (*b*) or unpaired (*u*). In the absence of any probing data, we assume each nucleotide has an equal probability of being paired or unpaired (*p*(*b*) = *p*(*u*) = 0.5). A reactivity measurement (*r_i_*) reduces the structural uncertainty, which can be quantified as

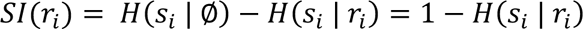

where *SI*(*r_i_*) denotes the structural information conveyed by reactivity *r_i_*, and *H*(*s_i_*) denotes the Shannon entropy of position *i*:

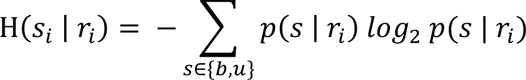

If *r_i_* conveys no information, *p*(*b* | *r_i_*) = *p*(*u* | *r_i_*) = 0.5 and *SI*(*r_i_*) = 0. Alternatively, if *r_i_* conveys perfect information, then (*b* | *r_i_*) = 1 or *p*(*u* | *r_i_*) = 1 and *SI*(*r_i_*) = 1. *p*(*b* | *r_i_*) and *p*(*u* | *r_i_*) are determined from the empirical reactivity distributions for paired and unpaired positions based on the known structure:

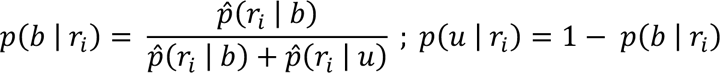

The distributions 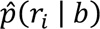 and 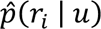 are estimated by fitting the paired and unpaired reactivity data for each nucleotide type to double-gamma mixture models. The expected structural information (ESI) is then obtained as the average over all nucleotides *n* in the molecule (excluding primer binding sites and other positions with low-quality data):

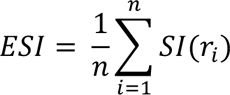

For computing ESI of DMS at only A and C nucleotides, *SI*(*r_i_*) of G and U nucleotides was set to 0.

### PAIR-MaP Analysis

We incorporated several minor updates to PairMapper analysis to maximize performance on four-base DMS datasets. *PairMapper* previously required nucleotide windows to have a minimum of 50 co-modification events to be considered for PAIR correlation analysis (8). We reevaluated this co-modification threshold across the range of 5 to 50, finding optimal performance with co-modification count cutoff of 10. We also updated the reactivity thresholds for primary and secondary PAIRs to 0.2 and 0.4, respectively.

Positive predictive value (ppv) and PAIR-MaP sensitivity (sens) were computed relative to accepted reference structures as previously described (8). RNA regions lacking DMS data were excluded from ppv and sens calculations.

### Structure Modeling

*Four-base DMS pseudo-energy parameterization:* We followed a previously described strategy (8, 47) to derive four-base-DMS-optimized pseudo-energy potentials for structure modeling in *RNAstructure* (v6.3) (48). Nucleotide-specific reactivity likelihood functions for paired and unpaired bases were fit using a double gamma mixture to normalized four-base DMS data collected on the cell-free *E. coli* 23S rRNA. These 23S rRNA derived parameters serve as universal folding parameters for all RNAs. Four-base DMS potentials only vary modestly from our previous nucleotide-specific DMS potentials (8), but more strongly penalize pairing of reactive G and U nucleotides. Fitted model parameters are provided in Supplemental Table S5. For structure modeling purposes, we replaced the DMSdist_nt.txt file in RNAstructure/data_tables/dists with our new four-base DMS-specific file. We plan to make these parameters available in future releases of *RNAstructure* via the *-fbDMS* flag.

*RNAstructure modeling:* Four-base DMS and SHAPE-directed modeling of RMRP, RNase P, and tmRNA was performed using iterative *ShapeKnots* folding to enable modeling of multiple pseudoknots (8, 49). *foldPK.py*, the automated script that facilitates this iterative folding strategy, is available for download at https://github.com/MustoeLab/StructureAnalysisTools. Folding of rRNAs was performed using *Fold* with the *-mfe* and *-md 600* options. Default SHAPE and four-base DMS parameters were used for all *Fold* and *ShapeKnots* modeling. For four-base DMS modeling, PAIR restraints were additionally passed using the *-x* option. Structure modeling was not possible for human 28S rRNA due to memory overflow errors in *RNAstructure*.

*Quantification of model accuracy:* The positive predictive value (ppv) and sensitivity (sens) of modeled structures were computed relative to accepted reference structures as previously described (8), using all Watson Crick and GU pairs allowing for one-position register shifts and ignoring singleton pairs. Accepted reference structures were obtained from refs (41-45). Modifications to tmRNA and RMRP structures were included as previously described (8).

## RESULTS

### Existing DMS-MaP strategies are unable to probe G structure in cells

To evaluate the ability of different MaP strategies to measure DMS modifications at all nucleotides, we generated MaP datasets from identical DMS-probed RNA inputs using published SSII (8), TGIRT (36), and Marathon (37) MaP protocols (Fig. 1A). We additionally evaluated an HIV-1 reverse transcriptase that was evolved to MaP N^1^-methyladenosine modifications (eHIV) (40), which has not been previously tested on DMS modified samples. DMS probing experiments were performed in duplicate on living HEK293 cells, under mildly alkaline buffer conditions that support multiple-hit DMS modification at all four nucleobases (see Methods) (8). An amplicon strategy was then used to obtain targeted DMS-MaP datasets for the Ribonuclease P (RNase P) and RnaseP RMP (RMRP) non-coding RNAs, which adopt well-defined, known structures.

**Figure 1.**
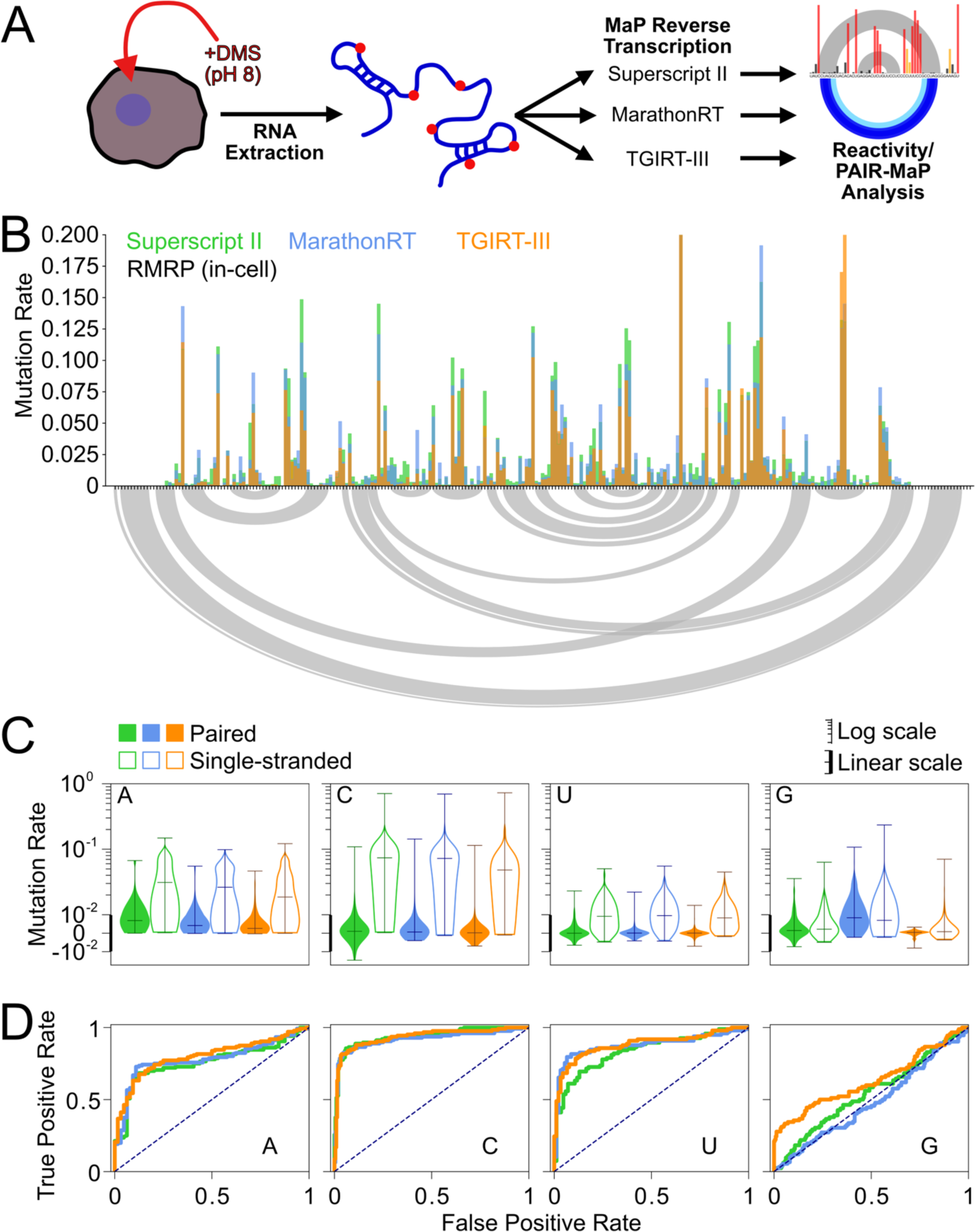
Comparison of different MaP protocols for measuring DMS modifications at all four RNA bases. **(A)** Experimental scheme for in-cell DMS probing, reverse transcription, and reactivity analysis using identical RNA inputs. **(B)** Background-subtracted DMS-MaP reactivity profiles measured for RMRP. Gray curves shown at bottom indicate known base pairing interactions. **(C)** DMS modification rates measured at each nucleotide combined across RMRP and RNase P. Background-subtracted mutation rates are shown for base-paired (filled) and single-stranded (open) bases. The y axis has a linear scale below <10^-2^ (indicated by thick axis) and logarithmic scale for values >10^-2^ (thin axis). **(D)** Receiver operator characteristic (ROC) curves quantifying ability of DMS reactivity to discriminate single-stranded versus base-paired nucleotides in RMRP and RNase P.

Consistent with prior studies (36, 37), analysis of untreated control samples revealed significant differences in the background error rates of different MaP protocols. While all protocols exhibit low median background rates (0.001, 0.001, and 5 × 10^-4^ for SSII, Marathon, and TGIRT respectively), SSII samples feature significantly more positions with high background rates (95^th^ percentiles of 0.02, 0.008, 0.005, respectively) (Fig. S1A). eHIV featured a 3-fold higher background rate than SSII (median 0.003; Fig. S1A), leading us to focus on the established MaP protocols for subsequent analyses.

Despite differences in background error profiles, background-subtracted DMS modification rates measured at adenosine (A), cytidine (C), and uridine (U) bases are broadly consistent across all enzymes (R > 0.9; Fig. 1B; Fig. S1B). As expected, Us are approximately 5-fold less reactive than A and Cs (Fig. 1C). U reactivities also vary more in SSII samples compared to Marathon and TGIRT, reflective of greater background noise in SSII samples. SSII and Marathon consistently measure higher modification rates compared to TGIRT (Fig. 1B, C), indicating differences in random nucleotide incorporation and enzyme drop-off. SSII also generates an increased fraction of indels compared to Marathon and TGIRT (26%, 2.8%, 7.4%, respectively; Fig. S1C). Nonetheless, each enzyme is similarly accurate at distinguishing single-stranded versus base-paired nucleotides (area under the receiver operating characteristic curve [AUROC] ≈ 0.79 for A; ≈ 0.93 for C; ≈ 0.83 for U; Fig. 1D; Table S1). These data corroborate our prior observation (8) that DMS is a highly specific probe of U pairing status and validate that all established MaP protocols reliably measure U modifications.

In contrast to A, C, and U nucleotides, all three MaP protocols exhibited minimal-to-no ability to measure G nucleotide structure (AUROC ≤ 0.6; Fig. 1D; Table S1). G mutation rates significantly increase upon DMS treatment, indicating that DMS is modifying G bases (Fig. 1C). However, these modifications occur at similar rates in both single-stranded and base-paired nucleotides. Surprisingly, G reactivity measurements vary 10-fold across MaP protocols, with SSII, Marathon, and TGIRT reporting median modification rates of 0.002, 0.007, and 8 × 10^-4^, respectively. Despite yielding the lowest overall modification rates, TGIRT does identify several highly reactive single-stranded Gs, but not with sufficient sensitivity to be useful. Thus, consistent with our prior studies (8), existing DMS probing protocols are unable to reliably probe G pairing status in cells.

### Mutational signature filtering enables robust DMS probing of G nucleotides

DMS is known to methylate G bases at two positions (2, 8). DMS predominantly modifies G at the N^7^ position with minimal dependence on Watson-Crick pairing status. At much lower rates, DMS can methylate single-stranded, deprotonated G nucleobases at the N^1^ position. While reverse-transcriptases are generally considered to be insensitive to N^7^-G modifications, we hypothesized that MaP may detect these modifications at low rates, convoluting any informative N^1^-G signal in DMS-MaP data. Further, we hypothesized that the differences in G modification rates measured by alternative MaP protocols reflect differences in N^7^-G detection efficiency.

To explore these hypotheses, we more closely analyzed the G mutational signatures yielded by each enzyme. DMS-MaP analysis traditionally considers all mutation types as conveying equivalent information. Strikingly, however, we observed major differences in the structural specificity of different mutation types. The elevated G modification rate measured by Marathon is driven by G→A substitutions, which occur at equivalent frequencies (median ≈ 0.005) in single-stranded and paired nucleotides (Fig. 2A; S2). SSII and TGIRT datasets similarly exhibit a surplus of structurally non-specific G→A substitutions, although at lower frequencies (median < 0.001). By contrast, DMS-dependent G→C and G→T substitutions occur almost exclusively in single-stranded nucleotides for all three enzymes (Fig. 2A). Thus, our DMS data are consistent with MaP measuring N^7^-G modifications specifically as G→A substitutions, whereas structurally informative N^1^-G modifications are decoded as G→C and G→T substitutions.

**Figure 2.**
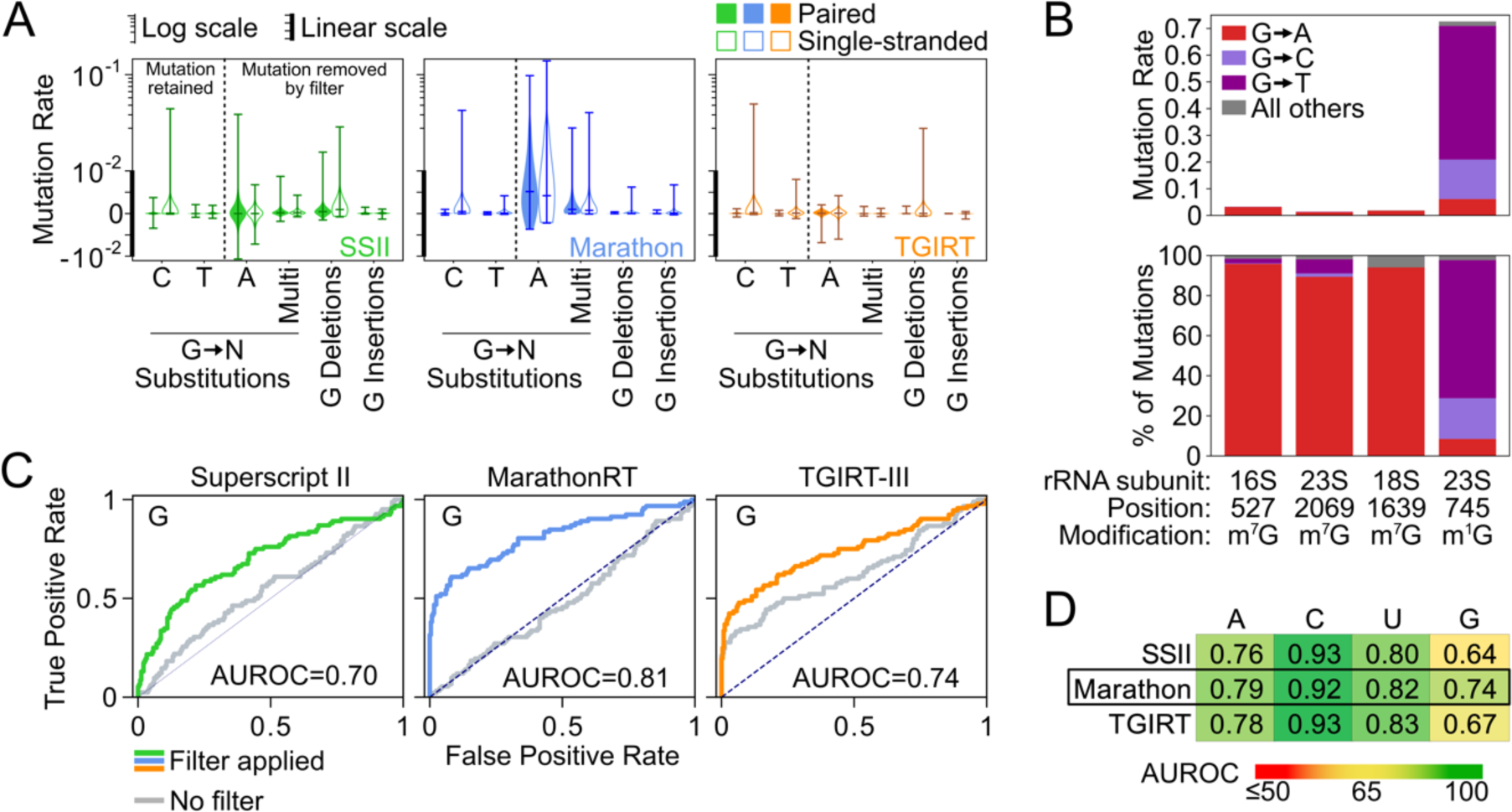
Mutation signature filtering enables high-fidelity DMS probing of G base-pairing status. **(A)** G-specific mutation spectrums for in-cell probed RMRP and RNase P generated by SSII (green), Marathon (blue), and TGIRT (orange). Rates are shown separately for base-paired (filled) and single-stranded (open) G nucleotides. The y axis has a linear scale below <10^-2^ (indicated by thick line) and logarithmic scale for values >10^-2^ (thin line). **(B)** Mutation rates (top) and percentage of detected mutations (bottom) measured by Marathon MaP for naturally-occurring N^1^-G and N^7^-G modifications in untreated *E. coli* and human rRNA. **(C)** ROC curves for mutation-signature-filtered G reactivities for in-cell probed RMRP and RNase P. Curves generated without mutation filtering for each enzyme are shown in gray. **(D)** Average AUROC across all probed RNAs quantifying the ability of mutation-signature-filtered DMS reactivities to discriminate pairing status at each nucleotide. The best performing enzyme, Marathon, is boxed. See Table S1 for the complete list of probed RNAs.

To validate this mutational signature, we used MaP to measure natural N^1^-G and N^7^-G modifications in untreated *E. coli* and human ribosomal RNAs (50, 51). Consistent with our DMS data, N^7^-G modifications are detected by Marathon with low efficiency (∼2%), but overwhelmingly as G→A substitutions (93% of mutations) (Fig. 2B). By comparison, the natural ribosomal N^1^-G modification is read out with high efficiency (65%) as a mixture of G→C and G→T substitutions (89% of mutations) (Fig. 2B). Similar results were also observed for SSII (not shown). Together, these data confirm that MaP decodes N^1^-G and N^7^-G modifications via distinct mutational signatures.

Leveraging this mutational signature, we implemented a refined bioinformatics pipeline within *ShapeMapper* (52) that filters out G→A substitutions and other uninformative mutation types in DMS probing data (see Methods). This refined pipeline resulted in dramatic improvements in the structural specificity of G DMS reactivity, with AUROC increasing from <0.6 to >0.7 for all three reverse-transcriptase enzymes (Fig. 2C). Benchmarking on an expanded panel of RNAs from human and *E. coli* cells confirmed that our pipeline enabled accurate DMS probing of G base-pairing status across diverse systems (Table S1). Marathon consistently yielded superior AUROC at G nucleotides (Fig. 2D, Table S1), leading us to select it as the optimal reverse transcriptase for DMS-MaP experiments. This improved strategy also reduced the importance of background mutation rate subtraction (Fig. S3), although the use of an untreated control still offers minor increases in probing accuracy. Overall, we conclude mutation-signature filtering enables robust DMS profiling of RNA structure at all four nucleotides, which we term four-base DMS-MaP.

### Appropriate buffering is essential for measuring U and G reactivities

The ability of DMS-MaP to measure structure-specific modifications at U and G requires transient deprotonation of N^1^-G and N^3^-U (8). We previously reported that bicine buffer (pH 8.0) is critical for robust DMS modification of G and U nucleotides *in vitro* (8). However, the extent to which extracellular buffering impacts DMS modification in cells is unclear. Cells work to maintain pH homeostasis, but changes in extracellular pH can affect intracellular pH, particularly on short time scales (53, 54). DMS treatment may also perturb the plasma membrane or induce stress responses that compromise internal pH control. We therefore investigated the impact of extracellular buffering by DMS probing *E. coli* and HEK293 cells at a variety of extracellular pHs: no supplemental buffer, which results in rapid acidification of the media (pH < 6); neutral pH 7.2 (sodium cacodylate buffer); and across the bicine buffering range (pH 7.7, pH 8.0, and pH 8.7). Both U and N^1^-G modification rates strongly depend on extracellular buffering, increasing ∼10-fold at pH 8 compared to unbuffered conditions (Fig. 3A, B). Interestingly, the reactivity rate of U and N^1^-G plateaus above pH 8, suggesting that cells buffer against major deviations from pH neutrality. The increase in G and U modification rate coincides with a significant increase in AUROC (from mean 0.76 to 0.91 for U, and 0.43 to 0.72 for G, respectively; Fig. 3C, D). By contrast, minimal changes in mutation rate or AUROC are observed at A or C nucleotides. The rate of N^7^-G modifications, measured by G→A substitutions, also minimally changes with pH (Fig. 3A, B). Thus, these data establish that extracellular pH — whether it is acidic due to DMS-induced acidification in the absence of buffer (10), neutral, or basic — modulates DMS reactivity in cells and emphasize that proper buffering is essential for four-base DMS probing. These data also provide further support for the deprotonation mechanism of DMS modification at N^1^-G and N^3^-U.

**Figure 3.**
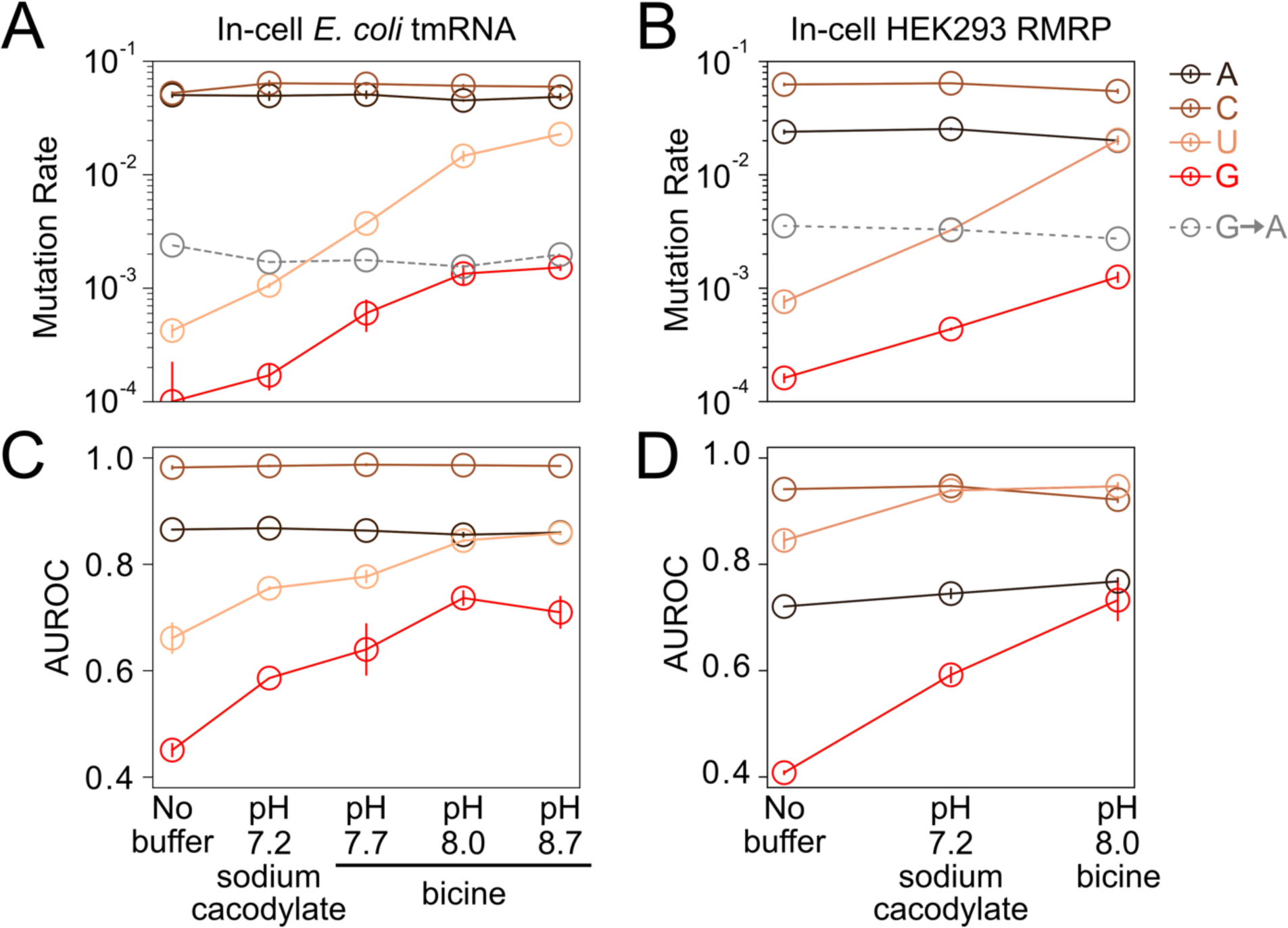
Four-base DMS-MaP depends strongly on extracellular buffering. **(A, B)** Mean DMS modification rates measured by Marathon MaP for single-stranded A,C,G, and U nucleobases for *E. coli* tmRNA and human RMRP probed in cells using different extracellular buffers. G→A substitutions, which are filtered out by mutation-signature-filtering, are shown in gray. **(C, D)** Corresponding AUROC values for *E. coli* tmRNA and human RMRP quantifying the ability of DMS reactivities to discriminate pairing status. All data points represent the mean of two independent biological replicates, with vertical bars indicating standard error. Lines are drawn between points to guide the eye.

### Four-base DMS data compare favorably to SHAPE data

To facilitate structural analysis, we implemented a strategy to correct for differences in DMS modification rates across bases and normalize reactivities to a common 0 to ∼1 scale, denoting unreactive and reactive nucleotides, respectively (Methods). Consistent with the AUROC analysis above, these normalized four-base DMS reactivities provide a precise map of RNA structure in cells with significantly increased resolution compared to traditional DMS data (Fig. 4A).

**Figure 4.**
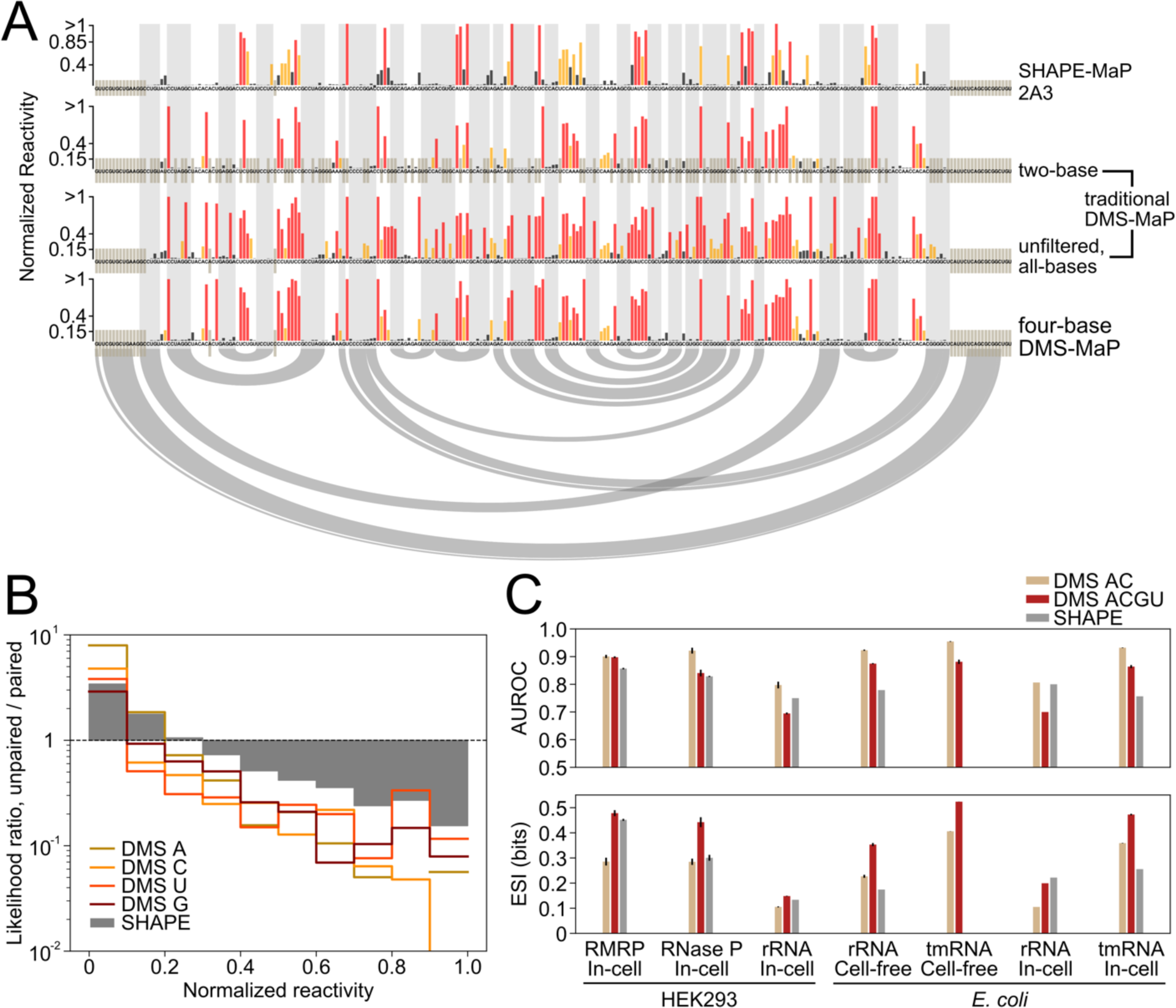
Four-base DMS-MaP conveys more structural information than other probing strategies. **(A)** Normalized reactivity profiles for in-cell probed RMRP given by SHAPE-MaP (top), traditional DMS-MaP (SSII; no mutation-signature filtering) considering only A and C nucleotides (upper middle) or all nucleotides (lower middle), or four-base DMS-MaP (bottom). Base pairing interactions in the accepted structure are shown using gray shading and arcs. **(B)** Unpaired/paired likelihood ratios for SHAPE and four-base DMS-MaP reactivities measured on cell-free probed *E. coli* 16S and 23S rRNA. A likelihood ratio of 1 indicates that a base has equal probability of being paired or unpaired given the measured reactivity. Likelihood ratios are shown for each nucleotide for four-base DMS-MaP, and are aggregated for all nucleotides for SHAPE. **(C)** Comparison of AUROC (top) and expected structural information (ESI; bottom) for four-base DMS-MaP and SHAPE-MaP across a diverse panel of RNAs. 2A3 SHAPE data for human rRNAs, and *E. coli* rRNAs and tmRNA were taken from ref. (34). AUROC and ESI values were also calculated for only A and C DMS reactivities, representing what is obtained by traditional two-base DMS probing experiments.

We sought to understand how four-base DMS reactivity data compare to SHAPE data, which represent the gold-standard for measuring structure at all four nucleotides (35). Prior *in vitro* studies have suggested that DMS and SHAPE data can convey similar amounts of structural information (8, 47), but direct comparisons of in-cell DMS and SHAPE data are lacking. Recently, 2A3 was introduced as an improved SHAPE reagent for in-cell structure probing, and high-quality 2A3 datasets are available for human and *E. coli* ribosomal RNAs (rRNAs) and *E. coli* tmRNA (34). We also collected our own 2A3 SHAPE-MaP data for RMRP and RNase P in HEK293 cells (Fig. S4). We note that we relied on published SHAPE protocols without pursuing further optimizations. Visual analysis indicates that SHAPE and four-base DMS reactivity profiles are qualitatively similar, although DMS provides a more binary measure of structure compared to a more continuous measure provided by SHAPE (Fig. 4A). Four-base DMS data also typically yield slightly higher AUROC than SHAPE data (Fig. 4C, S5).

Prompted by the differences observed between SHAPE and DMS data, we sought to develop a better analytic framework for evaluating the information conveyed by probing experiments. AUROC quantifies how well reactivity data perform as a binary classifier of a nucleobase being paired versus single-stranded, but this does not reflect how reactivity data are normally interpreted. For most applications, data are interpreted probabilistically: given a measured reactivity, what is the probability that a base is paired versus unpaired? These probabilities are represented by the paired/unpaired likelihood ratio function, with more extreme likelihoods indicating greater information content (55). The DMS likelihood ratio function is more extreme for A, C, and U nucleotides, indicating that DMS encodes more structural information at these nucleotides, whereas SHAPE encodes modestly more information at G (Fig. 4B). To quantify these differences, we developed a new metric termed expected structural information (ESI) that measures the total per-base information conveyed by a probing experiment (see Materials and Methods). ESI has units of bits and ranges between 0 (no information) to 1 (perfect specification of paired/unpaired status for all nucleotides) (Fig. S6). SHAPE experiments provide an average of 0.25 bits of ESI, whereas four-base DMS data convey an average of 0.35 bits (Fig. 4C), supporting that four-base DMS provides a more deterministic measure of base-pairing. Notably, two-base DMS data only convey an average of 0.23 bits of ESI, typically less than SHAPE (Fig. 4C). Together, these analyses indicate that four-base DMS experiments typically provide more structural information than other widely used probing strategies.

### Four-base DMS-MaP enables improved direct base-pair detection from single-molecule PAIR analysis

In addition to providing a per-nucleotide readout of base-pairing status, DMS probing data can be analyzed at the single-molecule level to identify nucleotides that undergo correlated modification. PAIR analysis (8) is a powerful strategy for detecting RNA duplexes from single-molecule probing data, and significantly improves the confidence and accuracy of structural analysis. We hypothesized that the improved specificity of four-base DMS-MaP would also benefit PAIR analysis. Indeed, PAIR analysis applied to four-base DMS-MaP data significantly outperformed analysis of traditional DMS-MaP data (SSII without mutation-signature filtering): the positive predictive value (ppv) of PAIR correlations increased from 0.73 to 0.84, and sensitivity (sens) increased from 0.18 to 0.31, respectively) (Fig. 5A,B; Fig. S7,S8; Table S2). Four-base DMS-MaP also enabled detection of PAIRs at lower read coverages (coverage of ∼300,000 vs ∼400,000 required for DMS-MaP (8)) (Fig. 5C).

**Figure 5.**
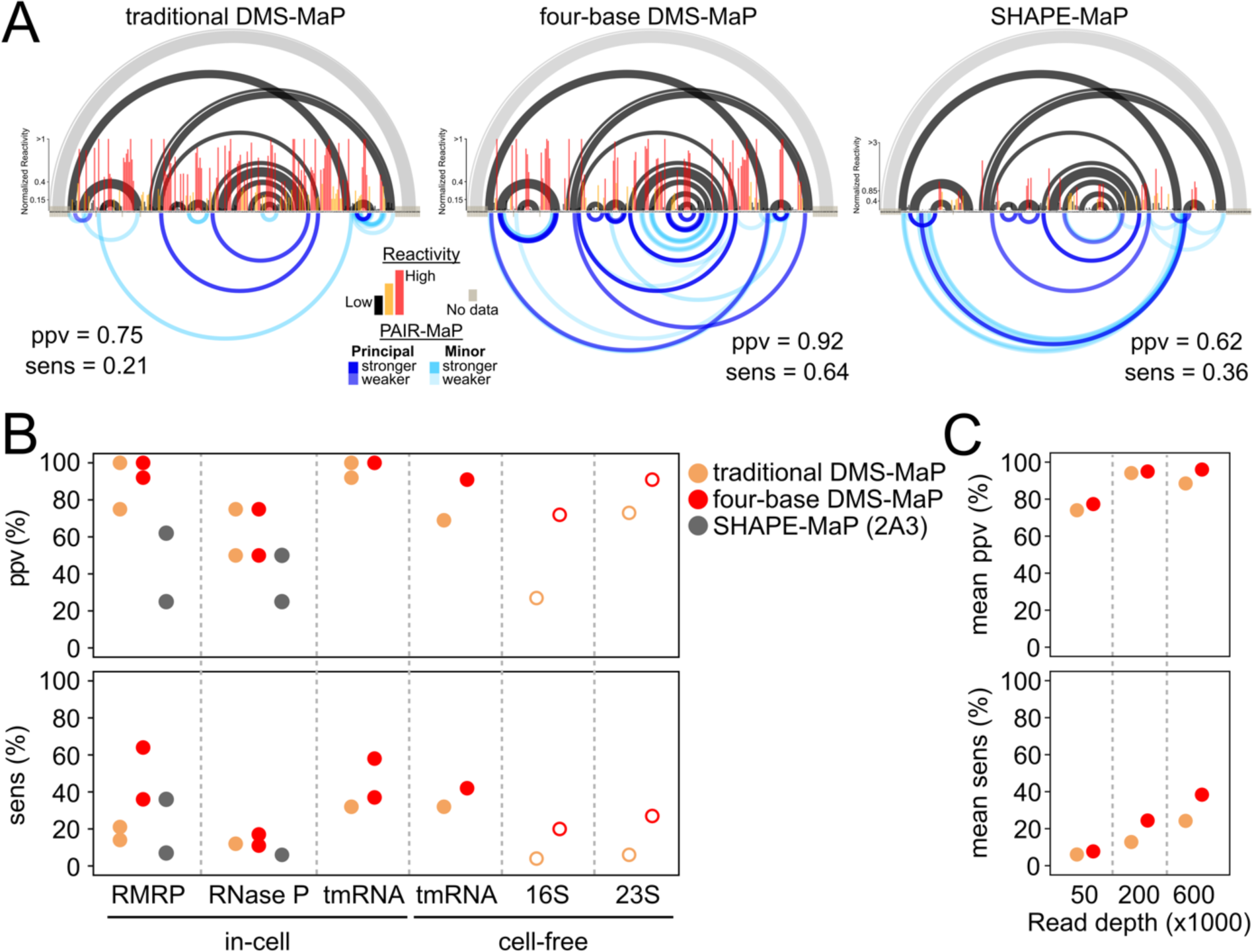
Four-base DMS probing improves single-molecule direct base-pair detection. **(A)** PAIR analysis performed on in-cell probed RMRP using traditional DMS-MaP (left), four-base DMS-MaP (center), or SHAPE-MaP (right) datasets. PAIRs are prioritized based on strength as principal and minor signals, shown in dark and light blue respectively. The accepted structure is shown at top. Long-range helices that are masked by primer binding sites, and thus lack PAIR-MaP data, are shown in light gray. PAIR positive predictive value (ppv) and sensitivity (sens) relative to the known structure are also indicated. **(B)** PAIR ppv and sens for traditional DMS-MaP (tan), four-base DMS-MaP (red), and SHAPE-MaP (gray). Data from two biological replicates are shown for in-cell RMRP, RNase P, and tmRNA. 16S and 23S rRNA data are shown in open circles and demonstrate reduced ppv and sens due to low sequencing coverage and likely misfolding of these RNAs under cell-free conditions. **(C)** Mean PAIR ppv and sens values for RMRP, RNase P, in-cell tmRNA, and cell-free tmRNA are shown for traditional DMS-MaP (tan) and four-base DMS-MaP (red) as a function of sequencing read depth. The indicated number of sequencing reads were unsampled without replacement from the larger datasets.

Surprisingly, PAIR analysis on traditional DMS-MaP data (SSII without mutation-signature filtering) performed worse than in prior studies (8). This reduced performance can be attributed to lower DMS modification rates (Fig. S9) and, for cell-free ribosomal RNA samples, significantly lower read depth coverage in our current experiments. We also note that many of the “false positive” PAIRs observed in cell-free rRNA samples likely correspond to “real” non-native interactions formed under these conditions (8, 56, 57). Interestingly, we also observed a 4-fold greater G→A substitution rate in our current DMS-MaP datasets compared to our prior experiments, suggestive of cryptic reverse-transcription differences that impact detection of N^7^-G modifications and reduce PAIR performance (Fig. S9). Mutation-signature filtering of SSII DMS-MaP data improved PAIR performance, but still underperformed Marathon DMS-MaP data (Fig. S7, S10).

We also performed PAIR analysis on four-base DMS-MaP datasets collected using TGIRT reverse-transcriptase. Compared to Marathon and SSII, TGIRT data gave significantly worse PAIR results (mean ppv = 0.57, sens = 0.20; Fig. S7, S10), presumably due to TGIRT measuring fewer modifications (Fig. 1, S10).

We additionally explored whether the 2A3 SHAPE reagent supports PAIR analysis. Historically, SHAPE reagents have been unable to achieve high enough modification rates for PAIR analysis. However, 2A3 addresses this limitation, modifying RNA at comparable rates to DMS, with lower modification of A and C compensated by higher modification at U and G (Fig. S4). Deeply sequenced human RMRP and RNase P in-cell 2A3 datasets both feature multiple PAIR correlation signals, but with lower sensitivity and specificity than four-base DMS-MaP (Fig. 5A, B). Thus, SHAPE experiments can enable PAIR detection, but further optimization is needed to make SHAPE-based PAIR analysis broadly useful.

### Four-base DMS-MaP enables improved RNA structure modeling

The end goal of many chemical probing studies is to translate probing data into an accurate model of RNA structure. Building on the improvements of four-base DMS-MaP, we developed new pseudo energy functions for incorporating four-base DMS reactivities as restraints during structure modeling with RNAstructure (Methods). Integrated structure modeling guided by four-base DMS reactivities and PAIR correlations facilitated accurate modeling of diverse, challenging RNA targets (Fig. 6). Notably, four-base DMS meets or exceeds the accuracy of SHAPE-directed structure modeling (Fig. 6). Four-base DMS data particularly benefits modeling accuracy for pseudoknot-containing tmRNA, RMRP, and Rnase P RNAs (Fig. 6), enabled by PAIR correlations unavailable to SHAPE. As an exception, SHAPE (2A3) data yields more accurate models for the in-cell 16S and 23S rRNAs, consistent with the superior ESI of 2A3 versus four-base DMS for these RNAs (Fig. 4C). Both four-base DMS and SHAPE data yielded similarly inaccurate models for in-cell human 18S rRNA; this inaccuracy arises from the extensive protein protections that reduce ESI of both reagents, as well as the poor thermodynamic stability of 18S rRNA (58). When the unrepresentative human and *E. coli* rRNAs are excluded, average ppv and sens of DMS-directed models increase to 0.93 and 0.92, respectively. In sum, four-base DMS-MaP supports best-in-class structure modeling accuracy and enables reconstruction of even challenging multi-pseudoknotted RNA structures.

**Figure 6.**
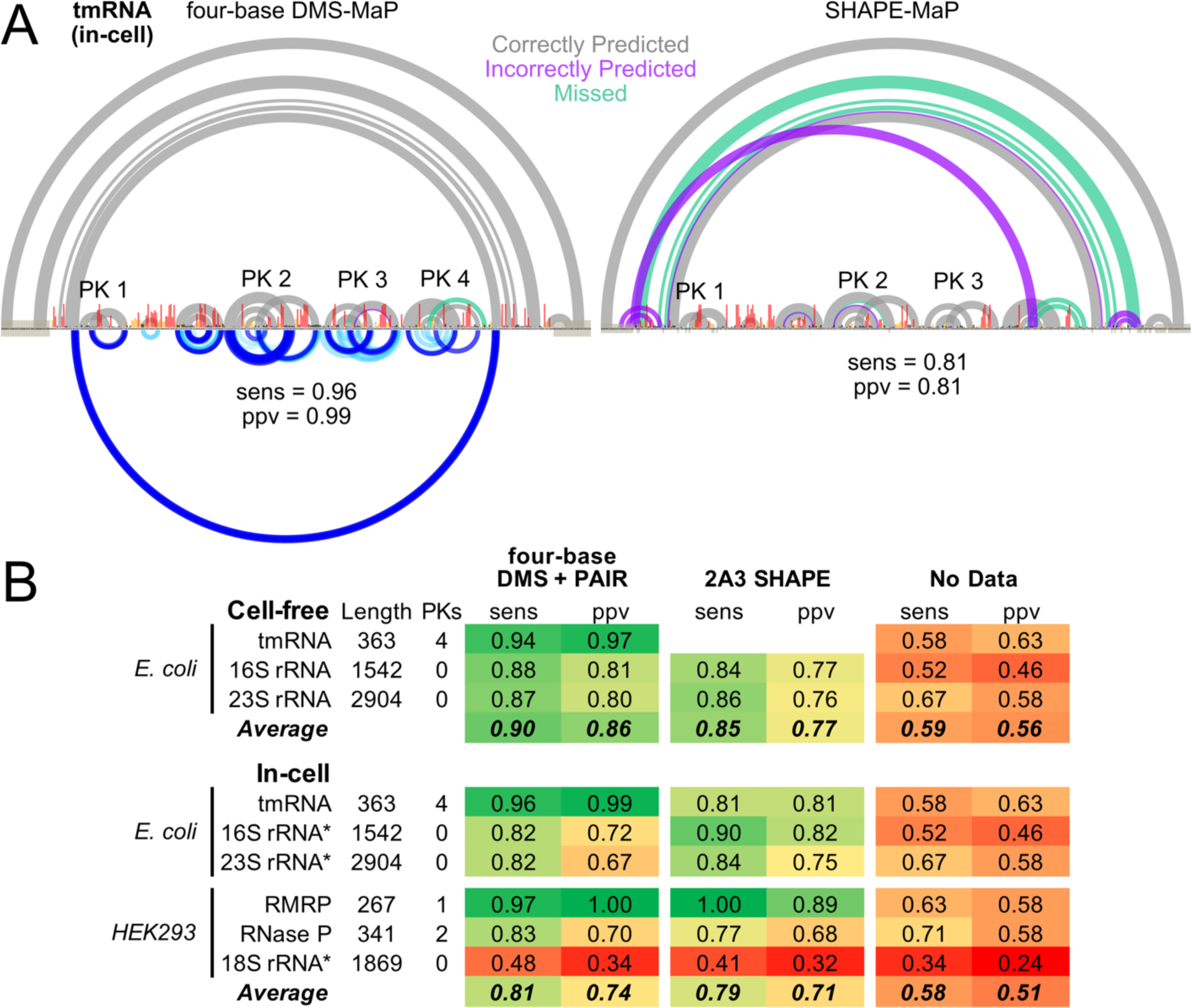
Four-base DMS-MaP enables improved RNA structure modeling. (**A**) *E. coli* tmRNA structure models obtained via *RNAstructure* modeling with four-base DMS and PAIR restraints (left) or SHAPE restraints (right) from in-cell experiments. True positive (gray), false positive (purple), and false-negative (green) predicted pairs are shown at top, and overall model ppv and sens are shown at bottom. For four-base DMS, measured PAIR correlations are also shown at bottom. Correctly modeled pseudoknots are labeled. **(B)** Structure modeling accuracy of four-base DMS-MaP and SHAPE-MaP. For in-cell RMRP, RNase P, and tmRNA, four-base DMS data from two biological replicates were pooled together. Asterisks (*) next to in-cell *E. coli* and HEK293 rRNAs indicate that four-base DMS structure modeling was done without PAIR restraints, which were not available due to insufficient sequencing depth. 2A3 SHAPE data for *E. coli* tmRNA and rRNAs, and human rRNAs were taken from ref. 34.

## DISCUSSION

DMS has long been a favored structure probing reagent and has played an essential role in enabling next-generation single-molecule probing analyses (4, 7). However, the inability of DMS to probe U and G nucleotides has been a critical limitation. In this work we introduced four-base DMS-MaP as a strategy for high-fidelity structure probing at all four nucleotides in living cells. Through rigorous benchmarking, we established that four-base DMS-MaP experiments typically convey more structural information than other available probing strategies and enhance single-molecule analysis, enabling accurate structure modeling of complex RNAs that challenge other methods. Four-base DMS-MaP experiments are straightforward to perform, requiring only minor changes to standard DMS probing protocols and bioinformatics pipelines. Thus, four-base DMS-MaP represents an “almost for free” upgrade offering improved resolution in both conventional per-nucleotide and single-molecule probing analysis.

The success of four-base DMS-MaP experiments depends on several subtle but collectively critical experimental parameters. Most importantly, our results emphasize the need for proper buffering, with G and U reactivity strictly dependent on pH (Fig. 3). Despite cells buffering against significant changes in intracellular pH, extracellular pH clearly modulates in-cell DMS reactivity. We also showed that different MaP protocols can impact data quality. SSII, Marathon, and TGIRT protocols performed similarly for per-nucleotide DMS reactivity analysis, although Marathon consistently performed the best at measuring N^1^-G modifications. However, the choice of MaP enzyme significantly impacted the success of single-molecule PAIR analyses, with Marathon consistently detecting more duplexes with fewer false positives than other MaP enzymes. TGIRT performed the worst at single-molecule PAIR analysis, likely because of a reduced ability to MaP through highly modified RNAs. We note that our analyses were limited to published MaP protocols, and speculate that optimization of MaP in the context of four-base DMS-MaP may yield even further improvements in data quality.

Four-base DMS-MaP is also built on the insight that MaP enzymes can simultaneously encode distinct types of chemical information via different mutational signatures. DMS modifies G nucleotides at two positions (59): The bulk of modifications occur at the N^7^ position, which do not report on Watson-Crick pairing, whereas only a minority of modifications occur at the informative N^1^ position. Consistent with other studies (60), our analysis indicates that Marathon (and to a lesser degree other reverse transcriptases) selectively decode N^7^-G modifications as G→A mismatches, allowing us to discriminate N^1^-G modifications and measure G pairing status with high fidelity. While not the focus of our current study, DMS N^7^-G modifications can provide information about RNA tertiary structure and G-quadruplexes (27, 60, 61), and further investigating the value of N^7^-G reactivity is a compelling area of future research. More generally, using mutation signatures to decode multiple coexisting modification signals represents a powerful paradigm for improving chemical probing analysis.

Our finding that four-base DMS-MaP typically conveys greater structural information than SHAPE-MaP is surprising. SHAPE chemistry holistically and unbiasedly measures nucleotide flexibility at the 2’ OH (62), but this holistic measure may come with the tradeoff of reduced specificity for Watson-Crick pairing compared to direct nucleobase probing by DMS. Reverse transcriptases may also decode 2’ OH SHAPE modifications with lower fidelity. Nevertheless, we emphasize that the performance gap between four-base DMS and SHAPE experiments is subtle, with both strategies performing well for most RNAs. SHAPE reagents also offer important benefits, including that they are generally less cytotoxic than DMS, are better suited for probing RNAs with modified bases, and, as noted above, holistically measure nucleotide flexibility. SHAPE reagents can further be used to probe all four nucleobases under single-hit reaction conditions, whereas four-base DMS probing of U, and especially G, nucleobases requires high overall modification rates. We also note that unlike for DMS, we made no attempt to optimize SHAPE-MaP. For example, SHAPE-specific data processing algorithms may enable improved PAIR analysis on SHAPE datasets. More generally, we believe that systematic efforts to improve MaP reverse-transcription and bioinformatics protocols will drive further increases in SHAPE probing resolution and accuracy.

Ultimately, our analysis demonstrates how improved chemical probing data can support further advances in RNA structure determination accuracy. The increased structural information provided by four-base DMS reactivities combined with PAIR correlation data is sufficient to guide in-cell structure modeling to >90% average accuracy for even very difficult targets such as tmRNA. Modeling accuracy is lower for rRNAs, but ribosomes are clearly exceptional cases with atypically high protein protections. Modeling accuracy is also reduced for human RNase P; follow-up analysis revealed that prediction accuracy was low even when using simulated “perfect” data (not shown), indicating that pseudoknot modeling algorithms remain imperfect. Combining four-base DMS-MaP with new statistical-learning strategies (63) represents one of several potential avenues for further improving modeling accuracy. Moving forward, we expect that focus will increasingly turn to the more difficult problem of modeling RNAs with heterogenous structures. Importantly, four-base DMS-MaP is fully compatible with emerging ensemble deconvolution analysis (11-14). We anticipate that the greater information provided by four-base DMS-MaP will help propel further advances in modeling and understanding complex RNA systems.

## DATA AVAILABILITY

Raw and processed probing data have been deposited at the GEO under accession number GSE225383. Analysis codes are available at https://github.com/MustoeLab and have been archived at 10.5281/zenodo.7808746.

## ACKNOWLEDGEMENTS

We thank D. Incarnato (U. Groningen) for sharing 2A3 data, and C. Weidmann (Michigan), M. Boerneke (UNC), and K. Weeks (UNC) for helpful comments. We thank L. Kearns (BCM), S. Busan (GSK), P. Irving (UNC), and K. Weeks (UNC) for ongoing support of *ShapeMapper*. We also thank H. Zhou (Boston College) and Y. Zhu and H. Zhai at the BCM Protein and Monoclonal Antibody Production Core for advice and help expressing and purifying the eHIV reverse transcriptase.

## FUNDING

A.M.M. is CPRIT Scholar in Cancer Research and a Beckman Young Investigator. The work was supported by the Cancer Prevention and Research Institute of Texas [RR190054] and the National Institutes of Health [R35 GM147010].

## CONFLICT OF INTEREST STATEMENT

As of April 2023, A.M.M. is an advisor to and holds equity in RNAConnect, Inc., which holds patent rights to MarathonRT. The research in this manuscript was conceived, funded, and executed independently of RNAConnect.

