## Supporting Information for "Mutation signature filtering enables high-fidelity RNA structure probing at all four nucleobases with DMS"

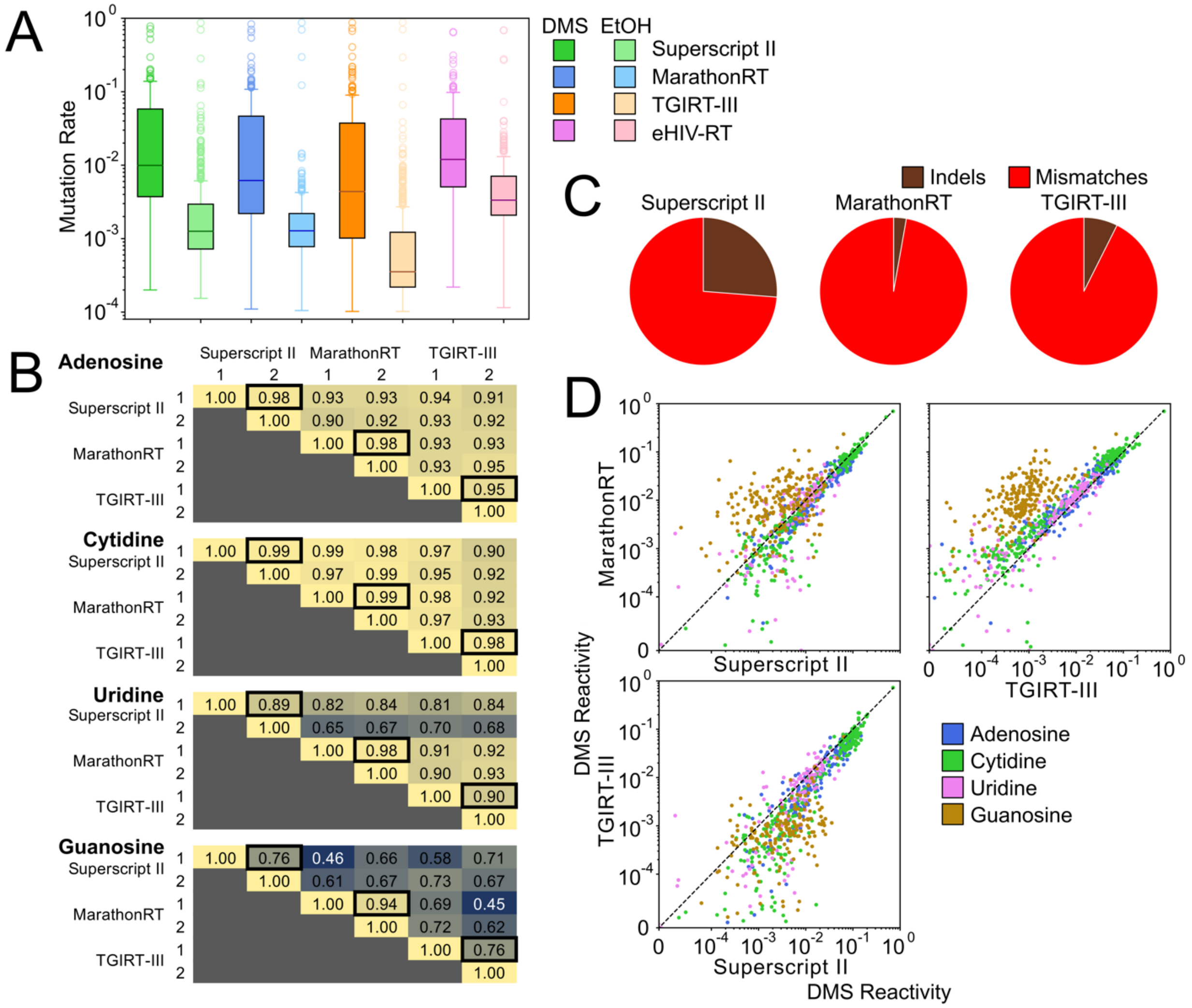


**Figure S1: (A)** Mutation rates of A and C nucleotides for in-cell DMS-treated (dark colors) and untreated (ethanol; light colors) RMRP and RNase P libraries generated by Superscript II (green), MarathonRT (blue), TGIRT-III (orange), or HIV-1 RT (violet). Mutation rates < $10^{-4}$ have been excluded. **(B)** Pearson's R correlations between background-subtracted DMS reactivities for two biological replicates measured using each reverse transcriptase. Data are combined for RMRP and RNase P. Black boxes denote correlations between biological replicates measured using the same enzyme. **(C)** Percentage of insertion-deletion (indels, brown) or mismatch mutations (red) generated by Superscript II, MarathonRT, and TGIRT-III for in-cell RMRP and RNase P. **(D)** Comparisons of background-subtracted DMS reactivities measured by Superscript II, MarathonRT, and TGIRT-III for RNase P and RMRP. Each point represents correlations for either A (blue), C (green), G (brown), or U (pink) nucleotides. The diagonal line represents identical DMS reactivity observed for both RTs.

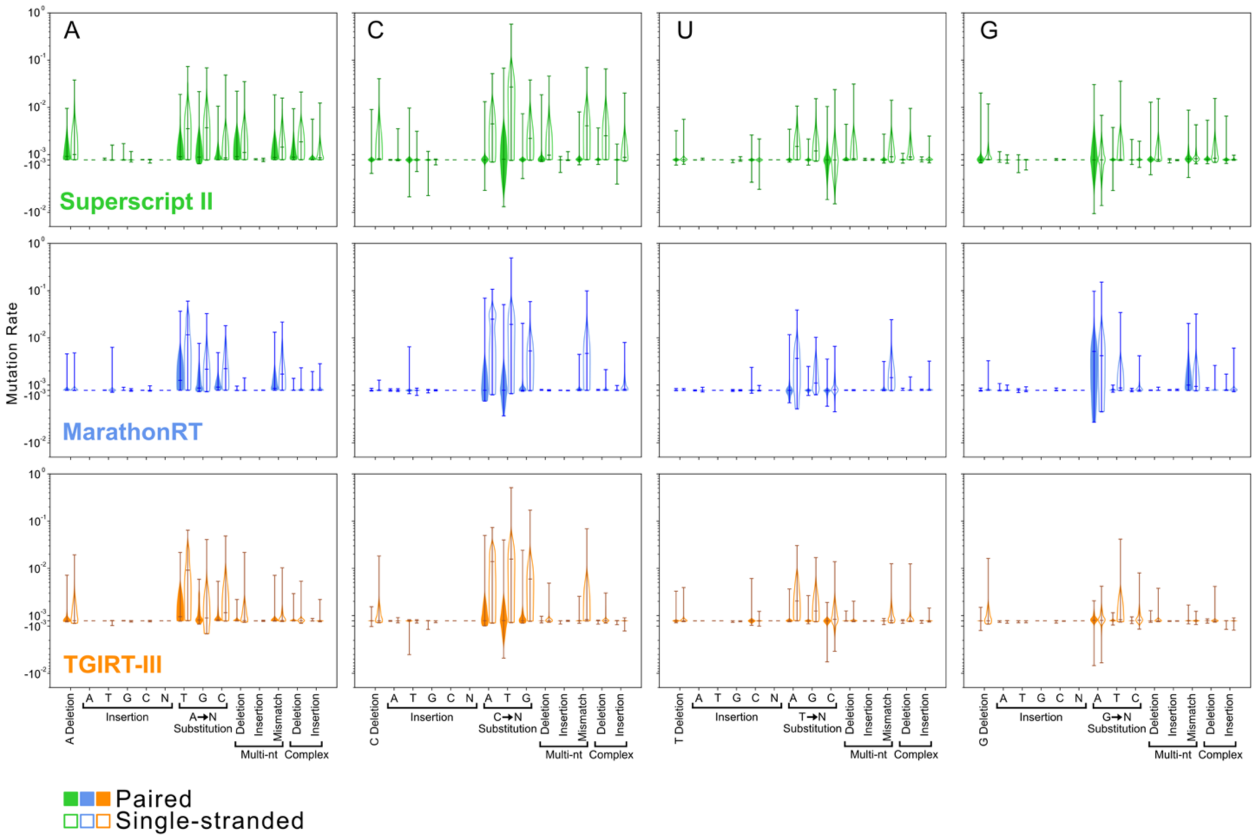


**Figure S2:** Mutation spectrums for in-cell DMS probing of combined RMRP and RNase P libraries generated using Superscript II (green), MarathonRT (blue), and TGIRT-III (orange). Background-subtracted mutation rates for base-paired (filled) and single-stranded (open) bases are shown for each mutation type.

| | | traditional DMS-MaP | | | | four-base DMS-MaP | | | |
|---|---|---|---|---|---|---|---|---|---|
| ***In-cell*** | Replicate | A | C | G | U | A | C | G | U |
| RMRP | 1 | -0.01 | 0.01 | 0.06 | 0.02 | -0.01 | 0.00 | 0.05 | 0.00 |
| | 2 | 0.00 | 0.01 | 0.06 | 0.01 | -0.01 | 0.01 | 0.04 | 0.02 |
| RNaseP | 1 | 0.01 | 0.06 | 0.06 | 0.10 | 0.01 | 0.00 | 0.02 | -0.06 |
| | 2 | 0.01 | 0.09 | 0.11 | 0.10 | 0.02 | -0.04 | 0.00 | -0.05 |
| tmRNA | 1 | 0.00 | 0.03 | 0.08 | 0.08 | 0.00 | 0.00 | 0.04 | 0.02 |
| | 2 | -0.01 | 0.02 | 0.08 | 0.09 | 0.00 | -0.01 | 0.03 | 0.02 |
| 16S rRNA | n/a | 0.02 | 0.06 | 0.02 | 0.08 | 0.02 | -0.02 | -0.02 | 0.03 |
| 23S rRNA | n/a | 0.01 | 0.05 | 0.07 | 0.06 | 0.01 | 0.00 | 0.00 | 0.01 |
| ***Cell-free*** | | | | | | | | | |
| tmRNA | n/a | 0.00 | 0.02 | 0.04 | 0.03 | 0.00 | 0.00 | 0.01 | 0.01 |
| 16S rRNA | 1 | 0.02 | 0.03 | 0.07 | 0.10 | 0.02 | 0.00 | 0.02 | 0.01 |
| | 2 | 0.00 | 0.00 | 0.00 | 0.00 | 0.00 | 0.00 | 0.00 | 0.00 |
| 23S rRNA | 1 | 0.02 | 0.02 | 0.09 | 0.07 | 0.01 | 0.01 | 0.01 | 0.01 |
| | 2 | 0.00 | 0.00 | 0.00 | 0.00 | 0.00 | 0.00 | 0.00 | 0.00 |
| ***Average*** | | ***0.01*** | ***0.03*** | ***0.06*** | ***0.06*** | ***0.01*** | ***0.00*** | ***0.02*** | ***0.00*** |

Background subtraction effect

- Increases AUROC
- No effect
- Decreases AUROC

**Figure S3:** The impact of background subtraction on accuracy of measured DMS reactivities. Shown is the difference in AUROC obtained when using background subtraction of an untreated control sample versus using only the DMS-modified sample. Analysis was done for both traditional DMS (left; SSII, no mutation signature filtering) and four-base DMS (right; Marathon, with mutation signature filtering). Blue indicates that background subtraction improves AUROC, white indicates no change, and red indicates a decrease in AUROC.

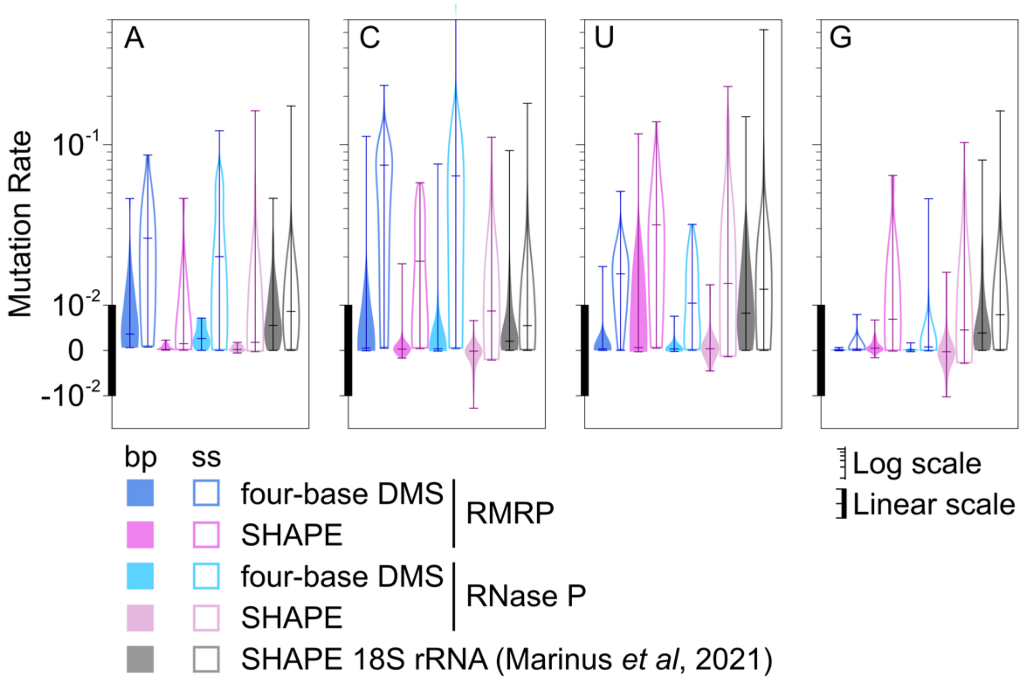


**Figure S4:** SHAPE 2A3 modification rates measured for in-cell probed RMRP and RNase P are comparable to four-base DMS and to prior 2A3 probing experiments. Background-subtracted modification rates are shown for four-base DMS and 2A3 probed RMRP and RNase P in HEK293 cells (current study), and for 18S rRNA data probed in HEK293 cells from Marinus *et al*, 2021 (1). Data are shown for base-paired (filled) and single-stranded (open) bases based on the accepted structures. The y axis has a linear scale below <$10^{-2}$ (indicated by thick line) and logarithmic scale for values >$10^{-2}$ (thin line).

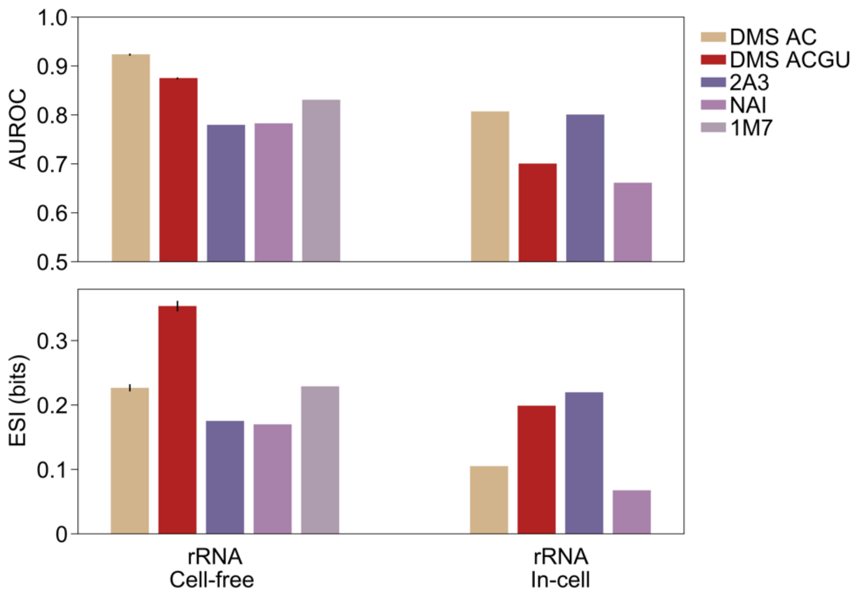


**Figure S5:** Comparison of AUROC and expected structural information (ESI) for probing *E. coli* rRNAs using four-base DMS and different SHAPE reagents (1,2). Error bars for DMS indicate standard deviation.

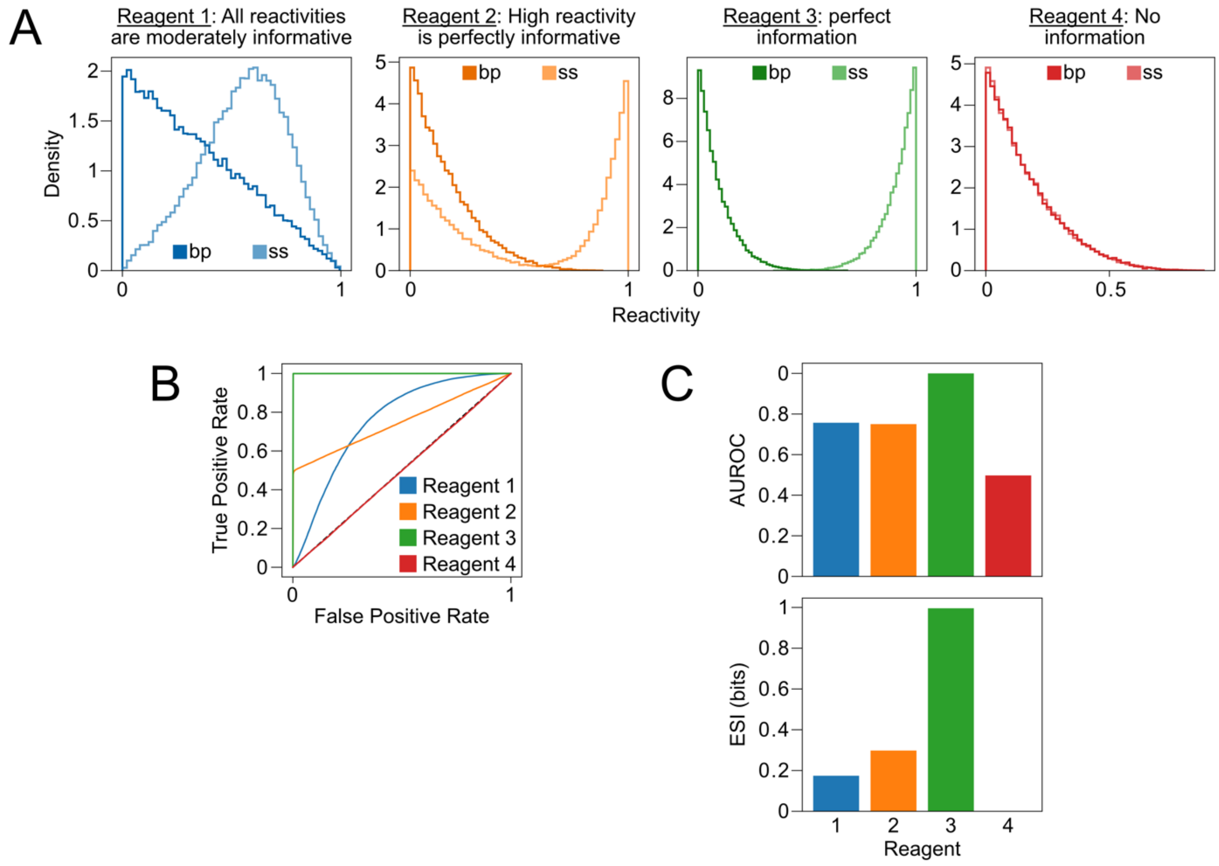


**Figure S6:** Expected structural information (ESI) provides a more complete measure of information yielded by a chemical probing experiment than AUROC. **(A)** Toy datasets illustrating different types of reactivity relationships for single-stranded and paired nucleotides. Reagent 1 preferentially modifies single-stranded nucleotides, but base-paired nucleotides are also modified at low-to-moderate rates. By comparison, Reagent 2 is extremely specific for single-stranded nucleotides and thus encodes more useful information for structure modeling purposes. Reagent 3 perfectly discriminates between paired and single-stranded nucleotides, with no overlap between the reactivity distributions, and Reagent 4 reacts indiscriminately with all nucleotides and thus encodes no information. (**B**) ROC curves generated from datasets in panel A. **(C)** Comparison of AUROC (top) and ESI (bottom) for datasets in panel A. Note that datasets 1 and 2 have identical AUROC values, but Reagent 2 yields higher ESI.

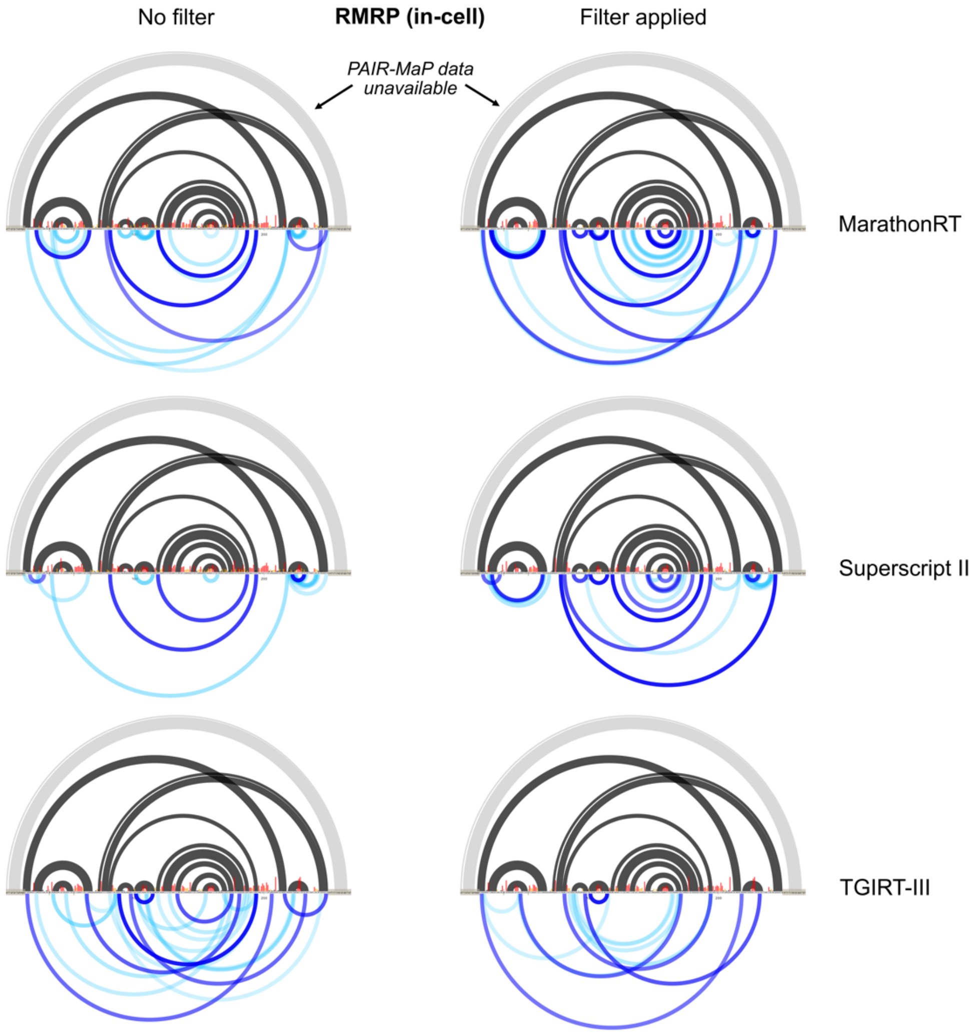


**Figure S7:** Mutation signature filtering improves PAIR direct base-pair detection for in-cell probed RMRP. PAIR-MaP data collected without (left) and with (right) mutation-signature filtering for MarathonRT (top), Superscript II (middle), and TGIRT-III (bottom). The accepted secondary structure is shown at top, with helices masked by primer-binding sites indicated in light gray. Principal (dark blue) and minor (dark blue) PAIR correlations are shown at bottom. Normalized per-nucleotide DMS reactivity (black, yellow, and red bars) are shown in the middle.

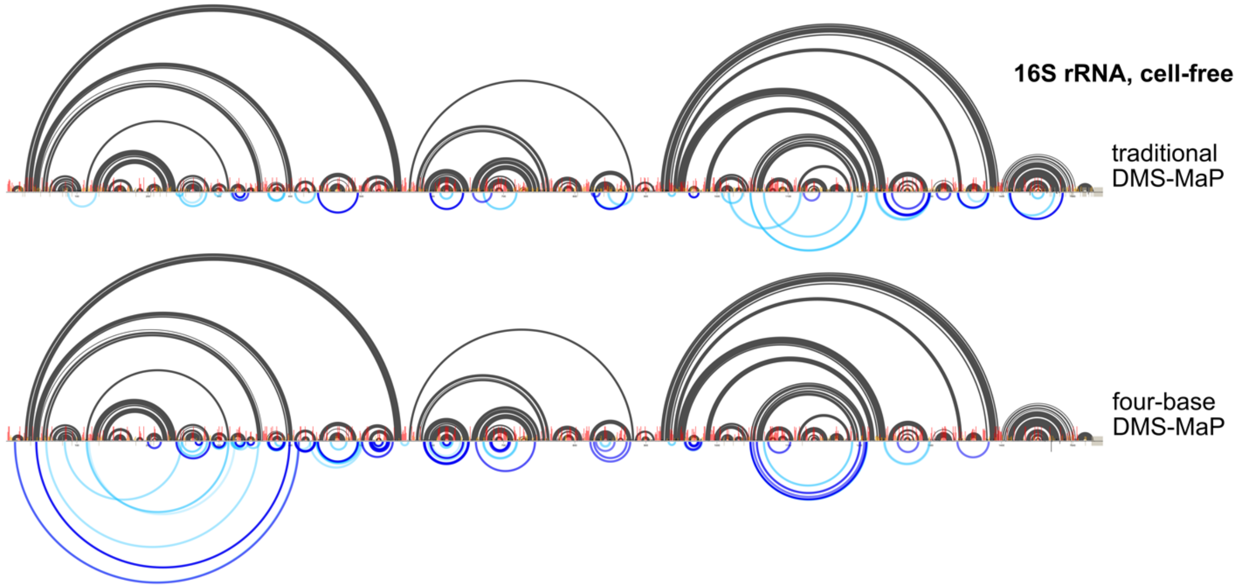


**Figure S8:** Mutation signature filtering improves PAIR direct base-pair detection for cell-free probed 16S rRNA. Shown are PAIR correlations measured in traditional DMS-MaP (SSII, no mutation signature filtering) and four-base DMS-MaP (Marathon, with mutation signature filtering) samples. The accepted structure is shown at top, and principal (dark blue) and minor (light blue) PAIR correlations shown at bottom. Normalized per-nucleotide DMS reactivities (black, yellow, and red bars) are shown in the middle.

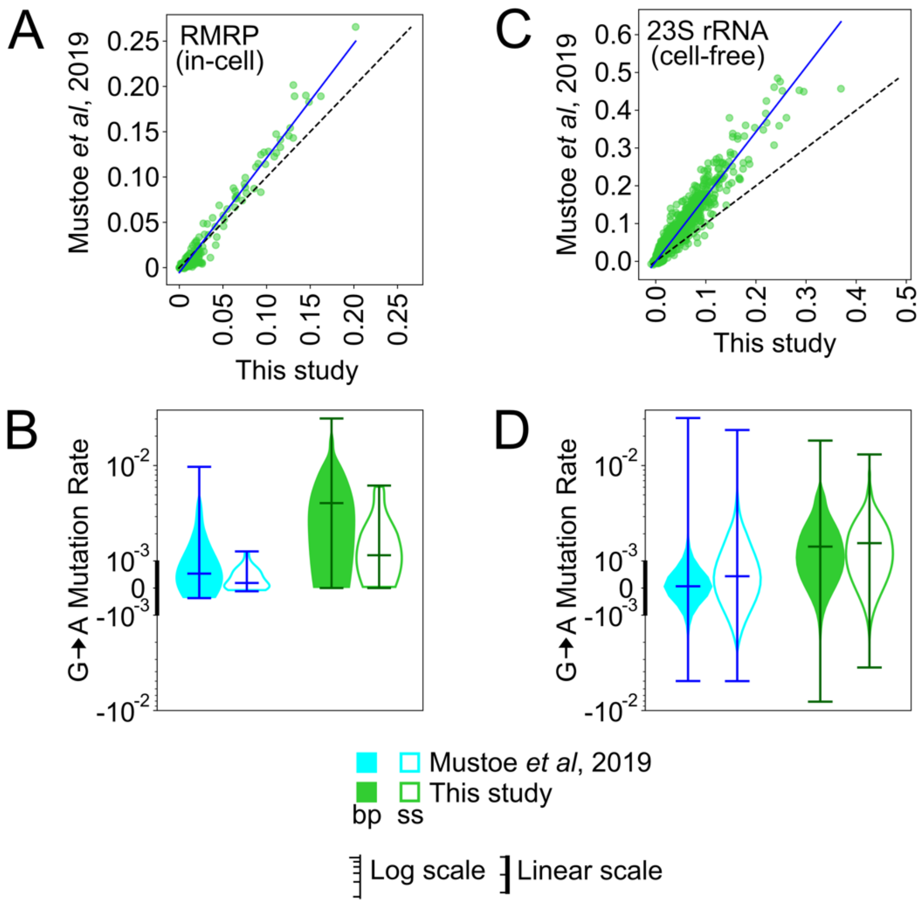


**Figure S9:** Comparison of SSII DMS-MaP libraries generated in this study and Mustoe *et al*, 2019 (3). **(A)** Correlation between background-subtracted DMS reactivities for in-cell probed RMRP in the two studies. The current study probed HEK293 cells, an adherent cell line, whereas Mustoe *et al*, 2019 probed Jurkat cells, a suspension cell line. **(B)** G-to-A mismatch mutation rates for RMRP from *Mustoe et al, 2019* (cyan) and this study (green), shown for base-paired (bp; filled) and single-stranded (ss; open) G nucleotides. **(C)** Correlation between background-subtracted DMS reactivities for cell-free *E. coli* 23S rRNA in the current study and reported in Mustoe et al. **(D)** G-to-A mismatch mutation rates for cell-free probed *E. coli* 23S from *Mustoe et al, 2019* (cyan) and this study (green), shown for base-paired (bp; filled) and single-stranded (ss; open) G nucleotides.

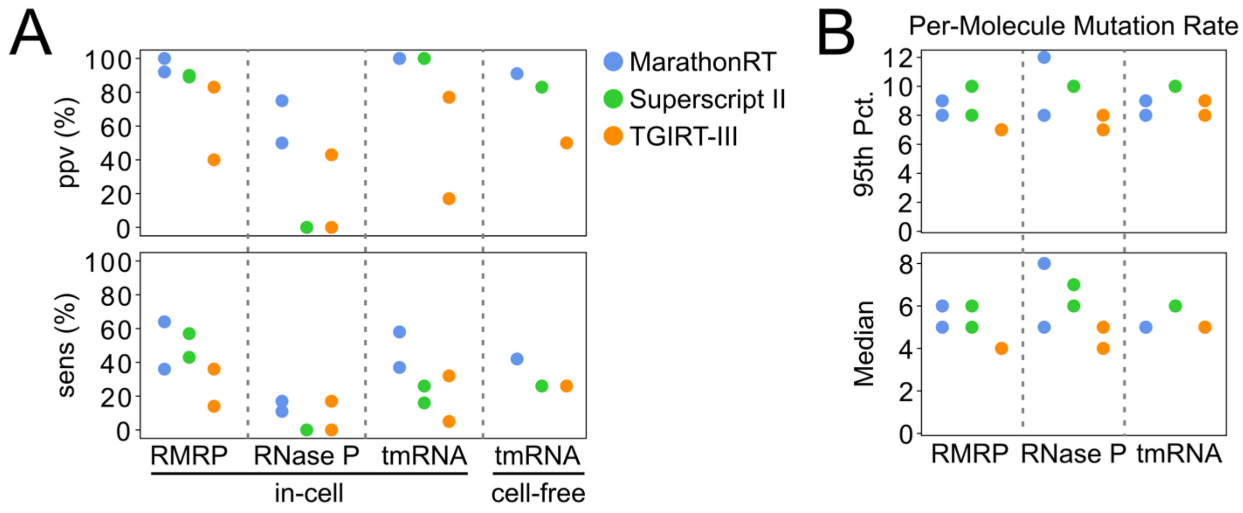


**Figure S10: (A)** PAIR ppv and sens for different MaP protocols with mutation signature filtering. Separate biological replicates for MarathonRT (four-base DMS-MaP; blue), Superscript II (green), and TGIRT-III (orange) are shown. Only principal PAIRs were included for ppv and sens calculations. **(B)** Median (bottom) and 95$^{th}$ percentile (top) mutation counts detected per sequencing read from in-cell probing experiments for MarathonRT (blue), Superscript II (green), and TGIRT-III (orange). Two biological replicates for in-cell human RMRP, RNase P, and *E. coli* tmRNA, are shown, and one replicate for cell-free *E. coli* tmRNA.

**Table S1:** Area under receiver operating characteristic curve (AUROC) values quantifying accuracy of DMS reactivities in discriminating base-paired versus single-stranded nucleotides.

**No Mutation Type Filtering**

| | | **Superscript II** | | | | **MarathonRT** | | | | **TGIRT-III** | | | |
|---|---|---|---|---|---|---|---|---|---|---|---|---|---|
| ***In-cell*** | Replicate | A | C | G | U | A | C | G | U | A | C | G | U |
| RMRP | 1 | 0.76 | 0.94 | 0.46 | 0.94 | 0.77 | 0.93 | 0.42 | 0.93 | 0.79 | 0.94 | 0.65 | 0.95 |
| | 2 | 0.73 | 0.93 | 0.53 | 0.88 | 0.77 | 0.92 | 0.40 | 0.95 | 0.77 | 0.93 | 0.63 | 0.96 |
| RNaseP | 1 | 0.78 | 0.95 | 0.58 | 0.81 | 0.80 | 0.93 | 0.45 | 0.80 | 0.83 | 0.96 | 0.69 | 0.87 |
| | 2 | 0.77 | 0.94 | 0.63 | 0.79 | 0.79 | 0.89 | 0.55 | 0.80 | 0.78 | 0.93 | 0.72 | 0.86 |
| tmRNA | 1 | 0.86 | 0.99 | 0.60 | 0.85 | 0.87 | 0.99 | 0.57 | 0.85 | 0.89 | 0.99 | 0.64 | 0.84 |
| | 2 | 0.85 | 0.99 | 0.60 | 0.84 | 0.87 | 0.99 | 0.56 | 0.85 | 0.88 | 0.99 | 0.62 | 0.81 |
| 16S rRNA | n/a | 0.62 | 0.82 | 0.51 | 0.69 | 0.64 | 0.85 | 0.47 | 0.68 | 0.66 | 0.86 | 0.54 | 0.64 |
| 23S rRNA | n/a | 0.60 | 0.83 | 0.45 | 0.68 | 0.61 | 0.87 | 0.43 | 0.66 | 0.61 | 0.83 | 0.52 | 0.68 |
| ***Cell-free*** | | | | | | | | | | | | | |
| tmRNA | n/a | 0.85 | 1.00 | 0.58 | 0.80 | 0.88 | 1.00 | 0.57 | 0.84 | 0.89 | 1.00 | 0.61 | 0.83 |
| 16S rRNA | 1 | 0.86 | 0.93 | 0.64 | 0.79 | 0.87 | 0.92 | 0.61 | 0.83 | n/a | n/a | n/a | n/a |
| | 2 | 0.83 | 0.91 | 0.67 | 0.81 | 0.85 | 0.91 | 0.67 | 0.83 | n/a | n/a | n/a | n/a |
| 23S rRNA | 1 | 0.80 | 0.92 | 0.62 | 0.80 | 0.82 | 0.92 | 0.65 | 0.86 | n/a | n/a | n/a | n/a |
| | 2 | 0.77 | 0.90 | 0.64 | 0.82 | 0.80 | 0.91 | 0.69 | 0.85 | n/a | n/a | n/a | n/a |
| ***Average*** | | 0.78 | 0.93 | 0.58 | 0.81 | 0.80 | 0.93 | 0.54 | 0.83 | 0.79 | 0.94 | 0.62 | 0.83 |

**Filtering Suboptimal Mutation Types**

| | | **Superscript II** | | | | **MarathonRT** | | | | **TGIRT-III** | | | |
|---|---|---|---|---|---|---|---|---|---|---|---|---|---|
| ***In-cell*** | Replicate | A | C | G | U | A | C | G | U | A | C | G | U |
| RMRP | 1 | 0.78 | 0.94 | 0.66 | 0.92 | 0.78 | 0.93 | 0.80 | 0.94 | 0.78 | 0.93 | 0.65 | 0.96 |
| | 2 | 0.72 | 0.93 | 0.60 | 0.86 | 0.77 | 0.92 | 0.76 | 0.96 | 0.77 | 0.92 | 0.65 | 0.97 |
| RNaseP | 1 | 0.83 | 0.94 | 0.69 | 0.82 | 0.79 | 0.93 | 0.72 | 0.80 | 0.82 | 0.95 | 0.81 | 0.88 |
| | 2 | 0.80 | 0.91 | 0.77 | 0.77 | 0.79 | 0.88 | 0.86 | 0.80 | 0.79 | 0.91 | 0.76 | 0.87 |
| tmRNA | 1 | 0.76 | 0.99 | 0.62 | 0.86 | 0.85 | 0.99 | 0.75 | 0.85 | 0.85 | 0.99 | 0.67 | 0.85 |
| | 2 | 0.76 | 0.99 | 0.57 | 0.86 | 0.85 | 0.98 | 0.74 | 0.84 | 0.84 | 0.99 | 0.76 | 0.83 |
| 16S rRNA | n/a | 0.64 | 0.85 | 0.52 | 0.67 | 0.64 | 0.85 | 0.56 | 0.67 | 0.66 | 0.87 | 0.56 | 0.63 |
| 23S rRNA | n/a | 0.62 | 0.83 | 0.50 | 0.66 | 0.62 | 0.86 | 0.54 | 0.66 | 0.62 | 0.84 | 0.55 | 0.67 |
| ***Cell-free*** | | | | | | | | | | | | | |
| tmRNA | n/a | 0.76 | 0.99 | 0.64 | 0.78 | 0.86 | 1.00 | 0.72 | 0.83 | 0.86 | 1.00 | 0.63 | 0.83 |
| 16S rRNA | 1 | 0.84 | 0.93 | 0.70 | 0.80 | 0.87 | 0.92 | 0.82 | 0.83 | n/a | n/a | n/a | n/a |
| | 2 | 0.80 | 0.91 | 0.71 | 0.81 | 0.85 | 0.91 | 0.80 | 0.83 | n/a | n/a | n/a | n/a |
| 23S rRNA | 1 | 0.79 | 0.92 | 0.70 | 0.79 | 0.82 | 0.93 | 0.79 | 0.85 | n/a | n/a | n/a | n/a |
| | 2 | 0.76 | 0.90 | 0.64 | 0.82 | 0.80 | 0.91 | 0.77 | 0.85 | n/a | n/a | n/a | n/a |
| ***Average*** | | 0.76 | 0.93 | 0.64 | 0.80 | 0.79 | 0.92 | 0.74 | 0.82 | 0.78 | 0.93 | 0.67 | 0.83 |

**Table S2:** Positive predictive value (ppv) and helix detection sensitivity (sens) values for principal PAIRs prior to and after filtering out uninformative mutation types. Single asterisks for sens indicates that values are colored on a different scale from ppv, reflecting that 50% sensitivity is typically sufficient to define global RNA architecture. Double asterisks (**) for rRNAs emphasize that these RNAs demonstrate reduced ppv and sens due to low sequencing coverage and misfolding of these RNAs under cell-free conditions.

**No Mutation Signature Filtering**

| | | MarathonRT | | Superscript II | | TGIRT-III | |
|---|---|---|---|---|---|---|---|
| ***DMS-MaP*** | Replicate | ppv | sens* | ppv | sens* | ppv | sens* |
| RMRP, in-cell | 1 | 0.80 | 0.29 | 0.75 | 0.21 | 0.38 | 0.14 |
| | 2 | 0.67 | 0.14 | 1.00 | 0.14 | 0.88 | 0.43 |
| RNase P, in-cell | 1 | 0 | 0 | 0.75 | 0.12 | 0.43 | 0.17 |
| | 2 | 0 | 0 | 0.50 | 0.12 | 0.50 | 0.06 |
| tmRNA, in-cell | 1 | 1.00 | 0.47 | 0.92 | 0.32 | 0.64 | 0.26 |
| | 2 | 1.00 | 0.37 | 1.00 | 0.32 | 0.33 | 0.11 |
| tmRNA, cell-free | – | 0.93 | 0.42 | 0.69 | 0.32 | 0.80 | 0.26 |
| 16S rRNA, cell-free** | – | 0.58 | 0.11 | 0.27 | 0.04 | n/a | n/a |
| 23S rRNA, cell-free** | – | 0.80 | 0.19 | 0.73 | 0.06 | n/a | n/a |
| ***Average*** | | ***0.64*** | ***0.22*** | ***0.73*** | ***0.18*** | ***0.57*** | ***0.20*** |
| ***SHAPE-MaP*** | | | | | | | |
| RMRP, in-cell | 1 | n/a | n/a | 0.62 | 0.36 | n/a | n/a |
| | 2 | n/a | n/a | 0.25 | 0.07 | n/a | n/a |
| RNase P, in-cell | 1 | n/a | n/a | 0.50 | 0.06 | n/a | n/a |
| | 2 | n/a | n/a | 0.25 | 0.06 | n/a | n/a |

**Mutation Signature Filtering Applied**

| | | MarathonRT | | Superscript II | | TGIRT-III | |
|---|---|---|---|---|---|---|---|
| ***DMS-MaP*** | Replicate | ppv | sens* | ppv | sens* | ppv | sens* |
| RMRP, in-cell | 1 | 0.92 | 0.64 | 0.90 | 0.57 | 0.40 | 0.14 |
| | 2 | 1.00 | 0.36 | 0.89 | 0.43 | 0.83 | 0.36 |
| RNase P, in-cell | 1 | 0.75 | 0.17 | 0 | 0 | 0.43 | 0.17 |
| | 2 | 0.50 | 0.11 | 0 | 0 | 0 | 0 |
| tmRNA, in-cell | 1 | 1.00 | 0.58 | 1.00 | 0.26 | 0.77 | 0.32 |
| | 2 | 1.00 | 0.37 | 1.00 | 0.16 | 0.17 | 0.05 |
| tmRNA, cell free | – | 0.91 | 0.42 | 0.83 | 0.26 | 0.50 | 0.26 |
| 16S rRNA, cell-free** | – | 0.72 | 0.20 | 0.47 | 0.06 | n/a | n/a |
| 23S rRNA, cell-free** | – | 0.91 | 0.27 | 0.70 | 0.08 | n/a | n/a |
| ***Average*** | | ***0.86*** | ***0.35*** | ***0.64*** | ***0.20*** | ***0.44*** | ***0.19*** |

**Table S3:** List of oligonucleotide primers used in this study.

| RNA | Sequence (5' → 3') |
|---|---|
| *E. coli* tmRNA | RT primer:<br>GAGCTGGCGGGAGTTGAA<br><br>Step1 forward primer:<br>CCCTACACGACGCTCTTCCGATCTNNNNNCTGGATTCGACGGGATTTGC<br><br>Step1 reverse primer:<br>GACTGGAGTTCAGACGTGTGCTCTTCCGATCTNNNNNGAGCTGGCGGGAGTTGAA |
| Human RMRP | RT primer:<br>ACAGCCGCGCTGAGA<br><br>Step1 forward primer:<br>GACTGGAGTTCAGACGTGTGCTCTTCCGATCTNNNNNGTTCGTGCTGAAGGC<br><br>Step1 reverse primer:<br>CCTACACGACGACGCTCTTCCGATCTNNNNNACAGCCGCGCTGAGA |
| Human RNase P | RT primer:<br>AATGGGCGGAGGAGAGTAG<br><br>Step1 forward primer:<br>GACTGGAGTTCAGACGTGTGCTCTTCCGATCTNNNNNAATGGGCGGAGGAGAGTAG<br><br>Step1 reverse primer:<br>CCCTACACGACGCTCTTCCGATCTNNNNNATAGGGCGGAGGGAAGC |

**Table S4:** Total mapped reads obtained for each RNA sample.

| | | MarathonRT | | | |
|---|---|---|---|---|---|
| | | Cell-Free | | In-Cell | |
| | *Replicate* | **DMS-Treated** | **Ethanol-Treated** | **DMS-Treated** | **Ethanol-Treated** |
| *E. coli* 16S rRNA | 1 | 3,526,021 | 1,026,421 | 731,679 | 223,846 |
| | 2 | 1,403,704 | 535,212 | – | – |
| *E. coli* 23S rRNA | 1 | 6,931,000 | 4,076,453 | 2,421,718 | 1,219,669 |
| | 2 | 2,224,121 | 2,337,332 | – | – |
| *E. coli* tmRNA | 1 | 1,994,886 | 591,971 | 1,107,481 | 376,937 |
| | 2 | – | – | 361,989 | 160,111 |
| *H. sapiens* 18S rRNA | – | – | – | 96,414 | 59,564 |
| *H. sapiens* 28S rRNA | – | – | – | 745,304 | 344,488 |
| *H. sapiens* RMRP | 1 | – | – | 2,911,223 | 860,331 |
| | 2 | – | – | 1,541,914 | 410,353 |
| *H. sapiens* RNase P | 1 | – | – | 1,516,737 | 517,163 |
| | 2 | – | – | 1,656,720 | 507,636 |
| | | **Superscript II** | | | |
| | | **Cell-Free** | | **In-Cell** | |
| | Replicate | **DMS-Treated** | **Ethanol-Treated** | **DMS-Treated** | **Ethanol-Treated** |
| *E. coli* 16S rRNA | 1 | 5,954,416 | 4,013,782 | 1,078,663 | 224,474 |
| | 2 | 2,683,104 | 244,617 | – | – |
| *E. coli* 23S rRNA | 1 | 12,795,673 | 12,444,352 | 2,925,324 | 1,010,444 |
| | 2 | 5,100,229 | 2,100,028 | – | – |
| *E. coli* tmRNA | 1 | 1,658,299 | 498,875 | 619,040 | 145,814 |
| | 2 | – | – | 461,657 | 118,384 |
| *H. sapiens* RMRP | 1 | – | – | 2,761,871 | 1,096,829 |
| | 2 | – | – | 4,137,983 | 368,258 |
| *H. sapiens* RNase P | 1 | – | – | 1,138,937 | 438,676 |
| | 2 | – | – | 1,569,445 | 500,705 |
| | | **TGIRT-III** | | | |
| | | **Cell-Free** | | **In-Cell** | |
| | Replicate | **DMS-Treated** | **Ethanol-Treated** | **DMS-Treated** | **Ethanol-Treated** |
| *E. coli* 16S rRNA | 1 | – | – | 169,327 | 13,266 |
| | 2 | – | – | – | – |
| *E. coli* 23S rRNA | 1 | – | – | 2,118,195 | 288,090 |
| | 2 | – | – | – | – |
| *E. coli* tmRNA | 1 | 17,710,657 | 3,684,928 | 595,455 | 174,735 |
| | 2 | – | – | 308,586 | 97,385 |
| *H. sapiens* RMRP | 1 | – | – | 1,956,280 | 1,693,372 |
| | 2 | – | – | 1,228,952 | 351,847 |
| *H. sapiens* RNase P | 1 | – | – | 1,607,636 | 621,267 |
| | 2 | – | – | 1,588,376 | 469,859 |
| | | **SHAPE (2A3)** | | | |
| | | | | **In-Cell** | |
| | | | | **2A3-Treated** | **DMSO-Treated** |
| *H. sapiens* RMRP | 1 | – | – | 1,946,040 | 1,045,658 |
| | 2 | – | – | 1,546,226 | 1,510,101 |
| *H. sapiens* RNase P | 1 | – | – | 1,478,196 | 1,203,896 |
| | 2 | – | – | 1,654,146 | 1,344,007 |

**Table S5:** Gamma mixture model parameters for *P*( *r* | *paired, X*) and *P*( *r* | *unpaired, X*) likelihood functions obtained from fits to cell-free 23S rRNA DMS reactivity histograms. $\pi_i$, $k_i$, $\theta_i$ denote the weight, shape, and scale parameters, respectively, for the two gamma components.

| | $\pi_1$ | $k_1$ | $\theta_1$ | $\pi_2$ | $k_2$ | $\theta_2$ |
|---|---|---|---|---|---|---|
| A paired | 0.5037219 | 1.223612 | 0.07226959 | 0.4962781 | 8.419402 | 0.00353477 |
| A unpaired | 0.9310149 | 2.70636 | 0.2190188 | 0.06898506 | 1.470389 | 0.03178099 |
| C paired | 0.61771 | 1.708509 | 0.00123037 | 0.38229 | 0.6454202 | 0.0784732 |
| C unpaired | 0.2393865 | 12.41265 | 0.08181064 | 0.7606135 | 0.7860596 | 0.7372148 |
| G paired | 0.9931628 | 0.745598 | 0.08874551 | 0.00683722 | 14.74002 | 0.08362662 |
| G unpaired | 0.9939772 | 0.646673 | 0.8534933 | 0.00602276 | 66.29347 | 0.1313313 |
| U paired | 0.9352064 | 0.7352539 | 0.0328988 | 0.06479357 | 0.5950865 | 0.3451798 |
| U unpaired | 0.2472543 | 4.976125 | 0.1967613 | 0.7527457 | 0.8044157 | 0.3844779 |
